# Maturation of Hippocampus-Medial Prefrontal Cortex Projections Defines a Pathway-Specific Sensitive Period for Cognitive Flexibility

**DOI:** 10.1101/2025.01.23.634573

**Authors:** Arely Cruz-Sanchez, Kathleen E. LaDouceur, Anusha Abdusalom, Helen Chasiotis, Radu Gugustea, Mehreen Inayat, Unza Mumtaz, Maryam Hasantash, Christoph Anacker, Maithe Arruda-Carvalho

**Affiliations:** University of Toronto Scarborough, Department of Psychology, Toronto, Ontario, Canada, M1C1A4; University of Toronto Scarborough, Department of Cell and Systems Biology, Toronto, Ontario, Canada, M1C1A4; Department of Psychiatry, Columbia University, and Research Foundation for Mental Hygiene, Inc. (RFMH), New York State Psychiatric Institute (NYSPI), New York, NY, 10032, USA

**Keywords:** Ventral CA1, intermediate CA1, electrophysiology, mice, cognitive flexibility, set shifting, optogenetics, chemogenetics

## Abstract

The septotemporal axis of the hippocampus separates it into domains with unique molecular, cellular, downstream connectivity and behavioral profiles, and yet very little is known about the ontogenesis of these highly specialized subcircuits. Here, we used viral tracing, optogenetic-assisted patch clamping, chemogenetics and behavior in mice to examine changes in domain-defined hippocampus efferent projections from postnatal day (P)10 to P60. We found distinct anatomical and synaptic developmental signatures in ventral and intermediate CA1 downstream connectivity, with unique contributions to the prelimbic and infralimbic subregions of the medial prefrontal cortex (mPFC). Juvenile inhibition of the ventral and intermediate CA1-mPFC pathways led to opposing modulation of adult cognitive flexibility, establishing a sex- and pathway-specific sensitive period preceding the stabilization of CA1-mPFC synaptic transmission. Our data elucidate domain- and target-defined postnatal maturation of hippocampus efferents, identifying juvenility as a CA1-mPFC sensitive period with crucial implications for early life influences on adult cognition.

## Introduction

The hippocampus (HPC) plays a key role in learning and memory^1,2^, anxiety-like behavior^3–5^ and stress^6^, with substantial evidence of distinct downstream pathway-specific contributions to these behaviors. Hippocampal projections to the lateral septum (LS) and basolateral amygdala (BLA) underlie fear memory processing and motivated behavior^7–9^, while projections to the thalamic nucleus reuniens (Re) and medial prefrontal cortex (mPFC) are involved in working spatial memory and episodic memory^10–12^. Hippocampal projections to the nucleus accumbens (NAc) and lateral hypothalamus (LH) have been linked to anxiety, social interaction and feeding behavior^9,13^. HPC topographical organization across the septotemporal axis^14–16^ distinguishes three HPC domains: dorsal, intermediate and ventral HPC, which diverge in their cellular composition^17–21^, efferent/afferent projection patterns^6,22,23^, population encoding^24,25^ and behavioral output^14,26–30^. This HPC tripartite differentiation is present from birth at both the gene expression and topographical afferent pattern levels^31,32^, but very little is known about how the postnatal maturation of the HPC and its downstream projections diverge across its domains.

Early life is also the time of peak onset for neurodevelopmental and psychiatric disorders^33–35^, which often feature deficits in the HPC and mPFC^36–43^ and HPC-mPFC connectivity^44–48^ in patients and animal models^38–40,44–53^. The HPC-mPFC pathway comprises unidirectional projections from the HPC intermediate and ventral domains to the mPFC^54–58^, and supports decision making, reward learning, fear, and anxiety-like behaviors^11,33,59–61^, all of which are (i) developmentally regulated^18,38^ and (ii) implicated in neurodevelopmental and psychiatric disorders^33,62–64^. Furthermore, similar to the sensory cortex^65–68^, both HPC^69–71^ and mPFC^51,72–76^ display heightened synaptic plasticity and behavioral features in early life consistent with critical periods, which may inform sensitivity to external influences such as stress^66,77–81,74^. Despite this convergence of factors in early life, it is unclear exactly how HPC-mPFC maturation may affect adult behavior and/or risk of neurodevelopmental and mental disorders.

Cognitive flexibility, or the ability to adjust behavior in response to changing conditions^82^, is an essential facet of adaptive behaviour necessary for survival. Studies in humans broadly implicate the prefrontal cortex in cognitive flexibility^82–84^, particularly in the case of extra-dimensional (ED) set shifting^85–87^, a finding replicated in primates^88,89^ and rodents^90–93^. Importantly, HPC lesions also feature deficits in ED set shifting in humans^87^. In rodents, developmental manipulations of ventral HPC impair ED set shifting^94,95^ and alter mPFC morphology^95^, implicating the HPC-mPFC pathway in ED set shifting. While this is consistent with other research tying HPC-PFC alterations to cognitive flexibility deficits in schizophrenia^96,97^, the precise contribution of HPC-mPFC pathways to higher-order cognitive flexibility remains unclear.

Despite important contributions to our understanding of HPC-mPFC oscillatory activity in neonatal to juvenile mice^46,55,98,99^, and changes in vHPC-mPFC plasticity from adolescence to adulthood^100^, a comprehensive survey of the postnatal development of HPC-mPFC synaptic connectivity, and how this developmental trajectory may affect adult cognitive function, is currently missing. To address this gap, we first compared the developmental profile of intermediate and ventral CA1 innervation to brain regions implicated in learning and memory (lateral septum (LS), basolateral amygdala (BLA), mPFC, thalamic nucleus reuniens (Re)), social, anxiety- and depression-like behavior (bed nucleus of the stria terminalis (BNST), lateral hypothalamus (LH), nucleus accumbens (NAc)). We then resolved the synaptic maturation of projections from ventral and intermediate CA1 to either prelimbic or infralimbic layer 5 mPFC pyramidal neurons using optogenetics-assisted patch clamping at postnatal day (P)10, P15, P21, P30 and P60. Finally, we examined the functional implications of these developmental changes by chronically inhibiting the vCA1- or iCA1-mPFC pathways during either juvenility (P15-P30) or adulthood (P60-P75) and testing animals a month later in a cognitive flexibility task. We found that juvenile inhibition of the vCA1-mPFC pathway led to a sex-specific deficit in ED set shifting which was not present when the same inhibition took place in adulthood. Surprisingly, juvenile (but not adult) inhibition of the iCA1-mPFC pathway improved adult ED set shifting in both sexes. Together, these data establish a timeline for the maturation of the HPC-mPFC pathway, highlighting septotemporal differences and target-defined asynchronies in anatomical and synaptic maturation guiding sensitivity to circuit disruption.

## Results

### Dorsal, intermediate and ventral CA1 neurons differ in their projection patterns in adult animals

To understand the developmental timeline of HPC efferent pathway innervation, we first compared the axonal terminal patterns of CA1 neurons from the dorsal, intermediate and ventral HPC domains in adult mice. For consistency with subsequent analyses, we infused the dCA1, iCA1 and vCA1 of P53 C57BL/6J mice with AAV1-ChR2-EYFP virus and processed the brains one week later for histology (Fig. S1A), an interval previously established by our group as sufficient for terminal transport^101^. Target regions were chosen based on their involvement in the regulation of memory and emotional behavior: NAc^102–104^, LH^4^, LS^105^, BNST^106^, BLA^4,107^, Re^108^ and mPFC^109–111^. Consistent with previous reports^14,15,23,112–115^, we saw domain-specific differences in CA1 efferent innervation, particularly between dorsal and intermediate/ventral CA1 projections (Fig. S1). We found strong projections from iCA1 and vCA1 to mPFC (Fig. 1SB), NAc core and shell (Fig. S1C), LS (Fig. S1D), BLA (Fig. S1E), LH (Fig. S1F) and Re (Fig. S1G) which were mostly absent in dCA1-infused brains. We noticed strong projections from vCA1 to the BNST (Fig. S1H), which were far weaker for iCA1 and absent for dCA1, as described in the literature116. While all HPC domains send projections to the LS (Fig. S1D), the pattern of LS innervations differs between dorsal and intermediate/ventral projections, with the former predominantly innervating dorsomedial LS, and the latter the ventromedial LS (Fig. S1D), as previously reported^117^. We then examined the mPFC, with a focus on its prelimbic (PL) and infralimbic (IL) subregions, given their contributions to memory and cognitive functions^12,30,96,118–122^. We saw strong projections from both iCA1 and vCA1 spanning PL and IL, with the strongest signal in IL and PL layer 6 (Fig. S1B). dCA1 axons in both PL and IL were only visible upon GFP signal amplification using immunofluorescence (Fig. S1B, left). Accordingly, we were unable to evoke terminal signals from this pathway when patching from either PL or IL pyramidal neurons, which prevented us from pursuing experiments exploring the synaptic maturation of dCA1-mPFC projections.

**Figure 1.**
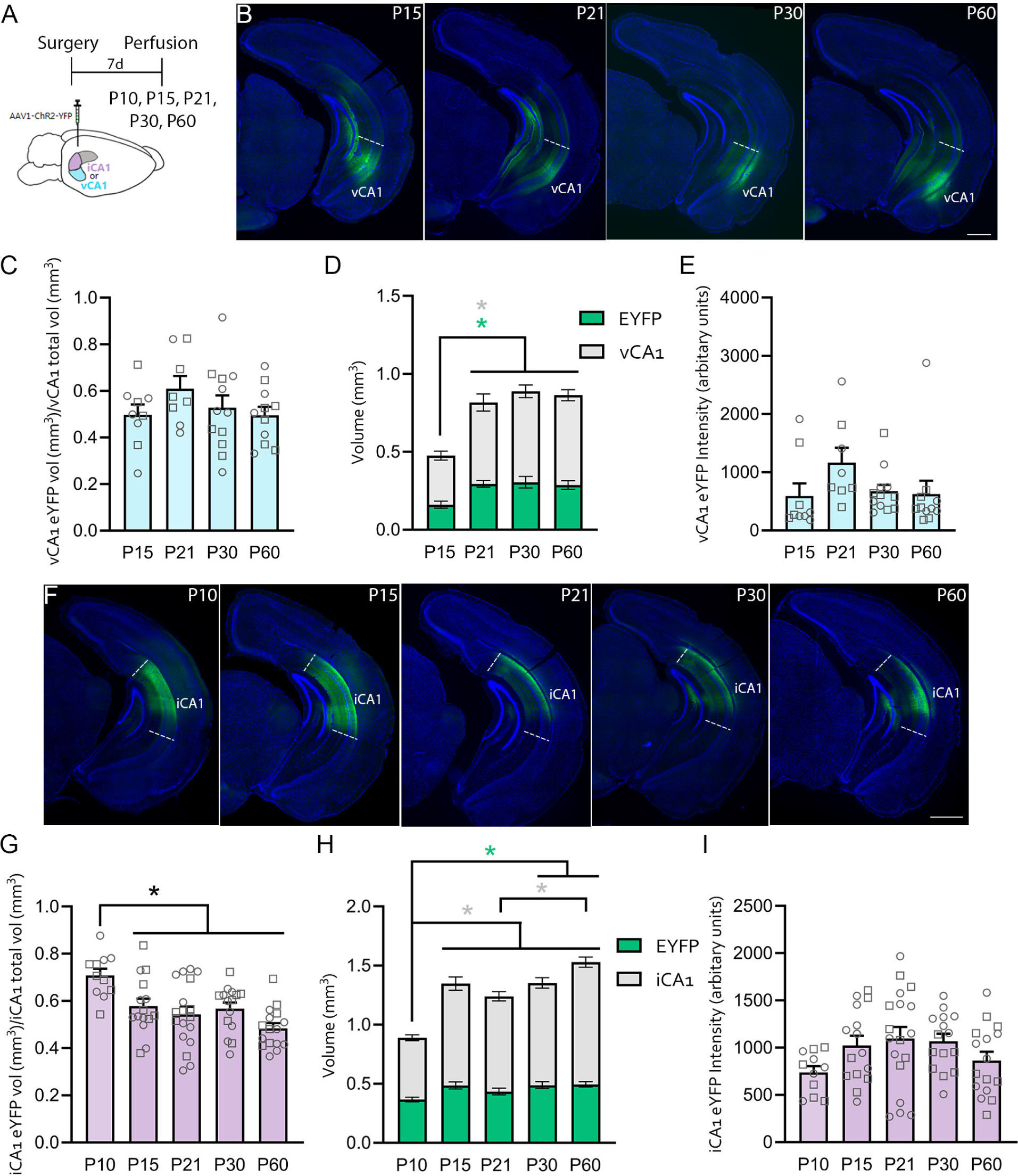
Targeting vCA1 and iCA1 across ages. **A.** C57BL/6J mice were injected with AAV-ChR2-YFP virus in either the vCA1 (**B-E**) or iCA1 (**F-I**) and perfused seven days later at P10, P15, P21, P30 or P60 for tissue and image processing. **B**. Sample images of AAV1-ChR2-EYFP infusion at the vCA1 in P15, P21, P30 and P60 mice. **C**. Estimated proportion of total vCA1 volume (mm^3^) with EYFP infection for each age group. **D**. Estimated total vCA1 volume (grey) and EYFP infected vCA1 volume (green) for all age groups. **E**. Average intensity of infected EYFP vCA1 area for each age group. **F**. Sample images of AAV1-ChR2-EYFP infusion at the iCA1 in P10, P15, P21, P30 and P60 mice. **G**. Estimated proportion of total iCA1 volume (mm^3^) with EYFP infection for each age group. **H**. Estimated total iCA1 volume (grey) and EYFP infected iCA1 volume (green) for all age groups. **I**. Average intensity of infected EYFP iCA1 area for each age group. Individual female datapoints are represented as circles, and males as squares. Scale bars: 500μm. **\*** p <0.05. Grey and green asterisks in **D** and **H** represent statistical significance in the total region volume and EYFP volume datasets, respectively. For full n values and break-down of all figures, refer to Table S1.

### Anatomical maturation of vCA1 and iCA1 projections across the lifespan

Given the lack of overlap in the terminal projection patterns from dCA1 compared to iCA1 and vCA1, we focused our efforts on comparing the maturation of iCA1 and vCA1 projections from infancy to adulthood. As ventral and intermediate CA1 projections overlap across most targets tested, we wanted to assess whether these subprojections would show equivalent maturation profiles, at both the anatomical and synaptic levels. First, we quantified CA1 targeting across ages to ensure robust viral transduction across ages and subprojections. Mice were injected with AAV1-ChR2-EYFP virus in either vCA1 or iCA1 and brains were processed for histology one week later at P10, P15, P21, P30 and P60 (Fig. 1A), corresponding to infancy, juvenility, late juvenility, adolescence and adulthood, respectively^123^. We observed that in mice younger than P15, vCA1 is very limited in total area compared to later ages, which precluded robust targeting of vCA1 (but not iCA1) at P10 without spread to CA3 and neighboring areas. Quantification of vCA1 infection area from P15 to P60 showed that the proportion of infected vCA1 area (i.e., ratio of vCA1 EYFP volume and total vCA1 volume) was equivalent across ages (Fig. 1B-C). The volume of infected vCA1 at P15 was lower than all older ages (Fig. 1D, green; one-way ANOVA p=0.0056; post hoc p<0.) to compensate for hippocampal growth during this period (Fig. 1D, grey; one-way ANOVA, p=0.00010; post hoc p< 0.013). There were no differences in vCA1 EYFP intensity (Fig. 1E) or sex differences in vCA1 volume across ages (Table S1).

Quantification of iCA1 infection showed a higher percentage of iCA1 targeted at P10, but not among other age groups (Fig. 1F-G; one-way ANOVA p=0.00010; post hoc p<0.036). This was likely driven by the significantly smaller volume of iCA1 at P10 (Fig. 1H, grey; one-way ANOVA p<0.00010; post hoc p<0.0022), despite a smaller iCA1 infected area at P10 compared to older ages (Fig. 1H, green; one-way ANOVA p=0.016; post hoc p<0.047). We found no age-dependent differences in EYFP iCA1 intensity (Fig. 1I) or sex differences in iCA1 volume across ages (Table S1).

#### vCA1 and iCA1 terminals reach target regions by P10

We next examined the age at which CA1 projections arrive to their target regions. vCA1 and iCA1 terminals were already present in all target regions by the earliest age tested of P10 (Fig. S2). To determine at what precise age CA1 axon terminals arrive at the mPFC, we infused RetroBeads into the mPFC and examined their transport to CA1 in infant and adult mice (Fig. S3A). We first confirmed that a 5-day incubation period was sufficient for retrograde transport from mPFC to CA1 in adult (8 weeks of age) mice (Fig. S3B). CA1 soma signal was present exclusively at the posterior/temporal HPC, at both the vCA1 and iCA1, but not at the anterior/septal (dorsal) HPC (Fig. S3B). We found soma signal in the CA1 of P10 brains following this same injection interval (Fig. S3C), consistent with our anterograde data (Fig. S2). We then tested the presence of CA1-mPFC projections at the earlier age of P5. There was no soma signal at the HPC of P5 brains (Fig. S3D), suggesting that CA1 axons are absent in the mPFC immediately after birth. As it is unclear how long RetroBeads can remain at a given target site without being uptaken by existing projections, these data conservatively suggest that CA1 axons arrive at the mPFC sometime between P0 and P10 in C57BL/6J mice.

#### Distinct target-defined maturation signatures in vCA1 and iCA1 terminal density

To characterize CA1 terminal density changes across ages, we quantified CA1 axon terminal fluorescent intensity in the NAc, LS, BLA, LH, Re, BNST and mPFC (PL and IL) (Fig. 2A-I; Fig. S4). vCA1 terminal density showed different maturation patterns among LS nuclei (Fig. 2A-C), showing no change in LSD (Fig. 2A; Fig. S4A) and LSI (Fig. 2C; Fig. S4A), but a higher density at P15 compared to P30 and P60 in the LSV (Fig. 2B; one-way ANOVA p=0.010; post hoc p=0.010; p=0.030, respectively). vCA1 terminal density remained stable across ages in the NAc (Fig. 2D-E; Fig. S4B). vCA1 terminal density in the BLA was higher at P15 and P60 compared to P21 (Fig. 2F; one-way ANOVA p=0.017; post hoc p<0.036; Fig. S4C), but remained stable in the Re (Fig. 2G; Fig. S4D). Interestingly, vCA1 terminal density in both BNST and LH showed a decrease between P15 and P21-30 (Fig. 2H-I; BNST: one-way ANOVA p= 0.0095; post hoc p= 0.044, p= 0.0077 respectively; LH: one-way ANOVA p=0.0068; post hoc p<0.05; Fig. S4E-F). We saw no sex differences in any of the regions examined (Table S1). These data suggest target-defined developmental changes in vCA1 efferent anatomical connectivity, with vCA1 terminal density in regions such as LSD, LSI, NAC and Re remaining stable from P15, while terminals in LSV, BLA, BNST and LH undergo further refinement across ages, emphasized by higher vCA1 terminal density at P15 in most of these regions.

**Figure 2.**
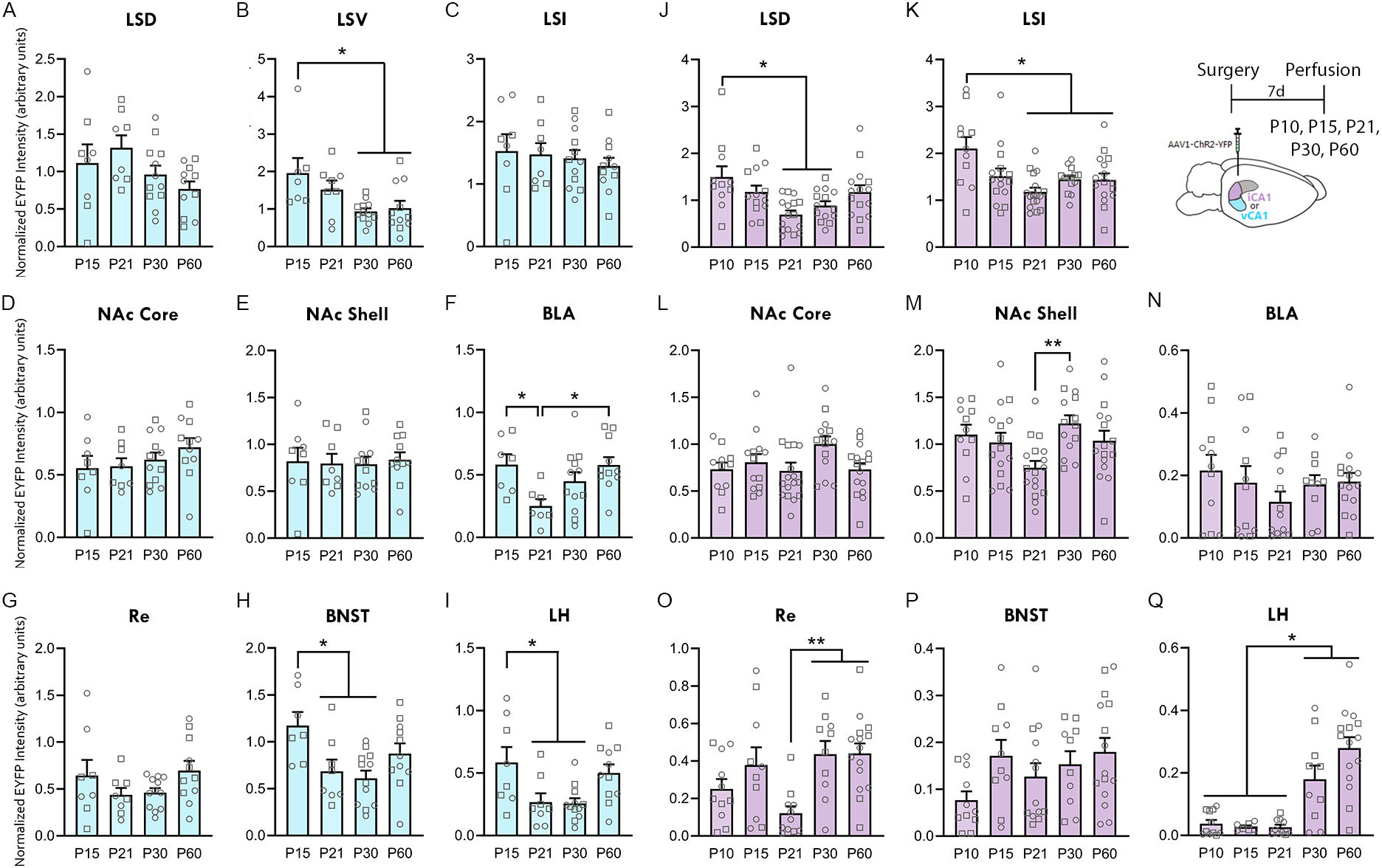
Pathway-specific developmental changes in vCA1 and iCA1 terminal density. C57BL/6J mice were injected with AAV-ChR2-YFP virus in the vCA1 (**A-I**) or the iCA1 (**J-Q**) and perfused seven days later at P10, P15, P21, P30 or P60 for tissue and image processing. **A-I**. Normalized average vCA1 terminal intensity across ages at the dorsal lateral septum (LSD; **A**), ventral lateral septum (LSV; **B**), intermediate lateral septum (LSI; **C**), nucleus accumbens (NAc) core (**D**) and shell (**E**); basolateral amygdala (BLA; **F**), thalamic nucleus reuniens (Re; **G**), bed nucleus of the stria terminalis (BNST; **H**), and lateral hypothalamus (LH; **I**). **J-Q**. Normalized average iCA1 terminal intensity across ages at the LSD (**J**), LSI (**K**), NAc core (**L**) and shell (**M**), BLA (**N**), Re (**O**), BNST (**P**), and LH (**Q**). Individual female datapoints are represented as circles, and males as squares. *p<0.05. **p<0.01.

We next conducted a similar analysis targeting the iCA1 (Fig. 2J-Q; Fig. S5). iCA1 terminal intensity in the LSD and LSI decreased with age, starting at P21 and ending at P30 in the LSD (Fig. 2J: one-way ANOVA p=0.0015; post hoc p<0.031; Fig. S5A), and at P21 in the LSI (Fig. 2K: one-way ANOVA p=0.0010; post hoc p<0.025; Fig. S5A). There were no iCA1 terminals in the LSV (Fig. S1D), indicating that vCA1 is the only source of LSV innervation across the CA1 septotemporal axis. Consistent with our results in the vCA1, iCA1 terminal intensity remained stable across ages in the NAc core (Fig. 2L; Fig. S5B), but increased between P21 and P30 in the NAc shell (Fig. 2M; one-way ANOVA p=0.010; post hoc p=0.0051; Fig. S5B). In further contrast to our vCA1 findings, there were no age-dependent changes in iCA1 terminal density in the BLA (Fig. 2N; Fig. S5C), and a surprising dip in iCA1 terminal density at P21 in the Re compared to P30-60 (Fig. 2O; one-way ANOVA p=0.0019; post hoc p<0.010; Fig. S5D) There was no change in iCA1 terminal density in the BNST (Fig. 2P; Fig. S5E), but an increase in the LH starting at P30 and continuing to P60 (Fig. 2Q one-way ANOVA p<0.00010; post hoc p<0.020; Fig. S5F). We found no sex differences in the regions analyzed (Table S1). These data emphasize a distinct pattern of terminal maturation between vCA1 and iCA1 projections, with regions that are predominantly stable across ages in one pathway (e.g., LSD, LSI, NAc shell and Re for vCA1) showing age-dependent changes in terminal density in the other pathway (e.g., iCA1), and vice-versa. A graphical summary of these findings is available in Fig. S6. Together, these data highlight domain- and target-specific time signatures of anatomical connectivity among HPC efferents.

We next quantified ventral and intermediate CA1 terminal density in the mPFC, focusing on PL and IL layers 2/3, 5 and 6 (Fig. 3A, H). We saw no differences in vCA1 terminal density from P15 to P60 in PL layers 2/3 (Fig. 3B), 5 (Fig. 3C) or 6 (Fig. 3D), nor across any layers in the IL (Fig. 3E-G). However, P15 females showed increased terminal density in PL and IL layer 6 compared to P15 males (PL: two-way ANOVA p=0.047; post hoc p=0.025; IL: two-way ANOVA p=0.037; post hoc p=0.039;Table S1). In contrast to the refinements seen in other vCA1 target regions, anatomical connectivity between vCA1 and the PL/IL remains stable from P15 onwards, with some differences between sexes.

**Figure 3.**
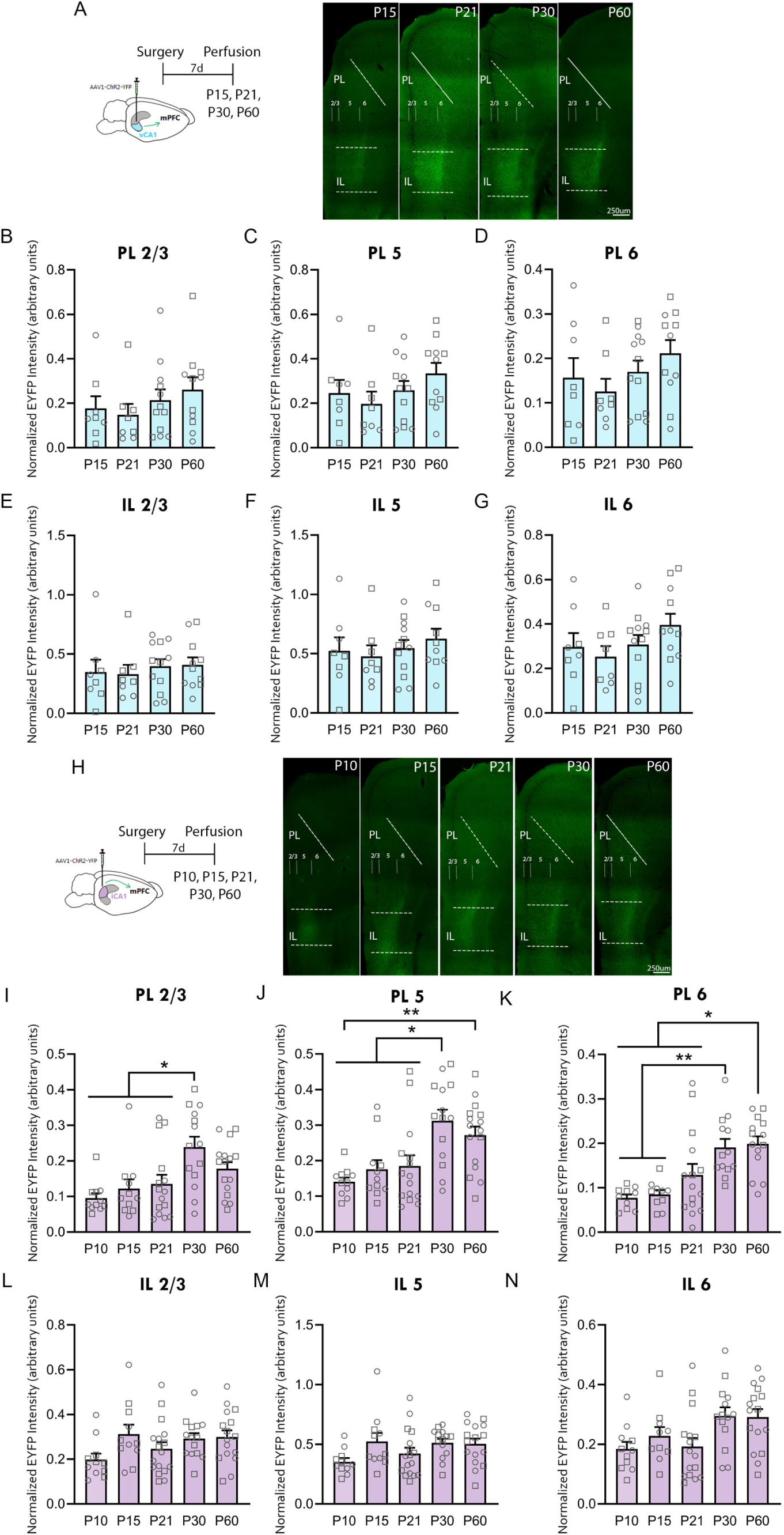
vCA1 terminal density is stable in PL and IL across postnatal development, but iCA1 terminal density increases in PL starting at adolescence. **A**. *Left*: Experimental design for the vCA1 experiments. C57BL/6J mice were injected with AAV-ChR2-YFP virus in the vCA1 and perfused seven days later at P15, P21, P30 or P60 for tissue and image processing. *Right*: Sample images of vCA1 projections present at the medial prefrontal cortex in P15, P21, P30 and P60 mice (numbers indicate cortical layers). **B-D**. Quantification of vCA1 terminal intensity across ages in PL layers 2/3 (**B**), 5 (**C**) and 6 (**D**)**. E-G**. Average vCA1 terminal intensity for all age groups at IL layers 2/3 (**E**), 5 (**F**) and 6 (**G**). **H**. *Left:* Experimental design for the iCA1 experiments. C57BL/6J mice were injected with AAV-ChR2-YFP virus in the iCA1 and perfused seven days later at P10, P15, P21, P30 or P60 for tissue and image processing. *Right*: Sample images of iCA1 projections present at the mPFC. **I-K**. Age-dependent differences in iCA1 terminal intensity in PL layers 2/3 (**I**), 5 (**J**) and 6 (**K**)**. L-N**. Average iCA1 terminal intensity for all age groups at IL layers 2/3 (**L**), 5 (**M**) and 6 (**N**). Individual female datapoints are represented as circles, and males as squares. Scale bar: 250μm. *p<0.05, ** p<0.01. PL = prelimbic cortex. IL = infralimbic cortex.

When we extended this analysis to iCA1 terminals (Fig. 3H-N), we found significant age-dependent increases in iCA1 terminals in all PL layers starting at P30 (Fig. 3I-K; Layer 2/3: one-way ANOVA p=0.00090; post hoc p<0.0020; layer 5: one-way ANOVA p<0.00010; post hoc p<0.010; layer 6: one-way ANOVA p<0.00010; post hoc p<0.050). These changes were not seen in IL (Fig. 3L-N). There were no sex differences in these datasets (Table S1). These data suggest asynchrony in the anatomical maturation of iCA1 terminals within the mPFC, with a delay in the stabilization of iCA1 innervation of PL relative to the IL, starting at adolescence. Furthermore, these divergent maturation trajectories point to significant septotemporal specialization in the anatomical maturation of iCA1 and vCA1 projections to mPFC, whereby only iCA1 projections to PL display age-dependent changes between infancy and adulthood.

### Maturation of CA1-PFC synaptic transmission

Dysfunction in the communication between the intermediate/ventral HPC and the mPFC is a prominent feature of cognitive deficits in rodent models of neurodevelopmental disorders^47^, and yet very little is known about the developmental trajectory of synaptic connectivity within this pathway^46,55,98,124^. To address this gap, we focused our investigation on the domain-specific synaptic maturation of CA1-mPFC pathways in C57BL/6J mice.

#### Maturation of spontaneous transmission in PL and IL layer 5 pyramidal neurons

To understand the context in which CA1 axons arrive and influence synaptic transmission in the mPFC, we first investigated the maturation of spontaneous excitatory and inhibitory synaptic transmission in layer 5 PL and IL pyramidal neurons at P10, P15, P21, P30 and P60. Starting with the PL, we saw an increase in the amplitude of spontaneous excitatory postsynaptic currents (sEPSCs) between P10-P21 to P30-P60 (Fig. 4A-B; one-way ANOVA p<0.00010; post hoc p<0.014), but no change in sEPSC frequency (Fig. 4C) or tau decay time (Fig. 4D). When examining PL spontaneous inhibition, we found a similar increase in the amplitude of spontaneous inhibitory postsynaptic currents (sIPSCs) between P10-21 and P30-P60 (Fig. 4E: one-way ANOVA p<0.00010; post hoc p<0.046) and an earlier increase in sIPSC frequency starting at P21 (Fig. 4F; one-way ANOVA p <0.00010; post hoc p<0.031), but no change in sIPSC decay time (Fig. 4G). A within-cell comparison of sEPSC and sIPSC PL frequencies showed a shift toward increased relative sIPSC frequency at P30 and P60 (Fig. 4H; paired t-test p<0.019), which is replicated in the increase in the spontaneous I-E frequency difference at P30 and P60 (Fig. 4I; one-way ANOVA p=0.0020; post hoc p<0.035). We saw sex differences in PL sEPSC and sIPSC amplitude (higher in P30 females and P60 males), sEPSC frequency (higher in males at P21 and P30) and sIPSC frequency (higher in males at P30 and P60), sIPSC tau decay time (higher in P60 females), as well as spontaneous I-E frequency differences, with P21 males and P60 females showing greater differences (Table S1). These data point to significant pre- and postsynaptic changes in spontaneous inhibitory transmission in PL layer 5 pyramidal neurons between P15-21 and P30, including a shift toward increased spontaneous inhibition that is likely reflective of maturation of local interneurons^125^.

**Figure 4.**
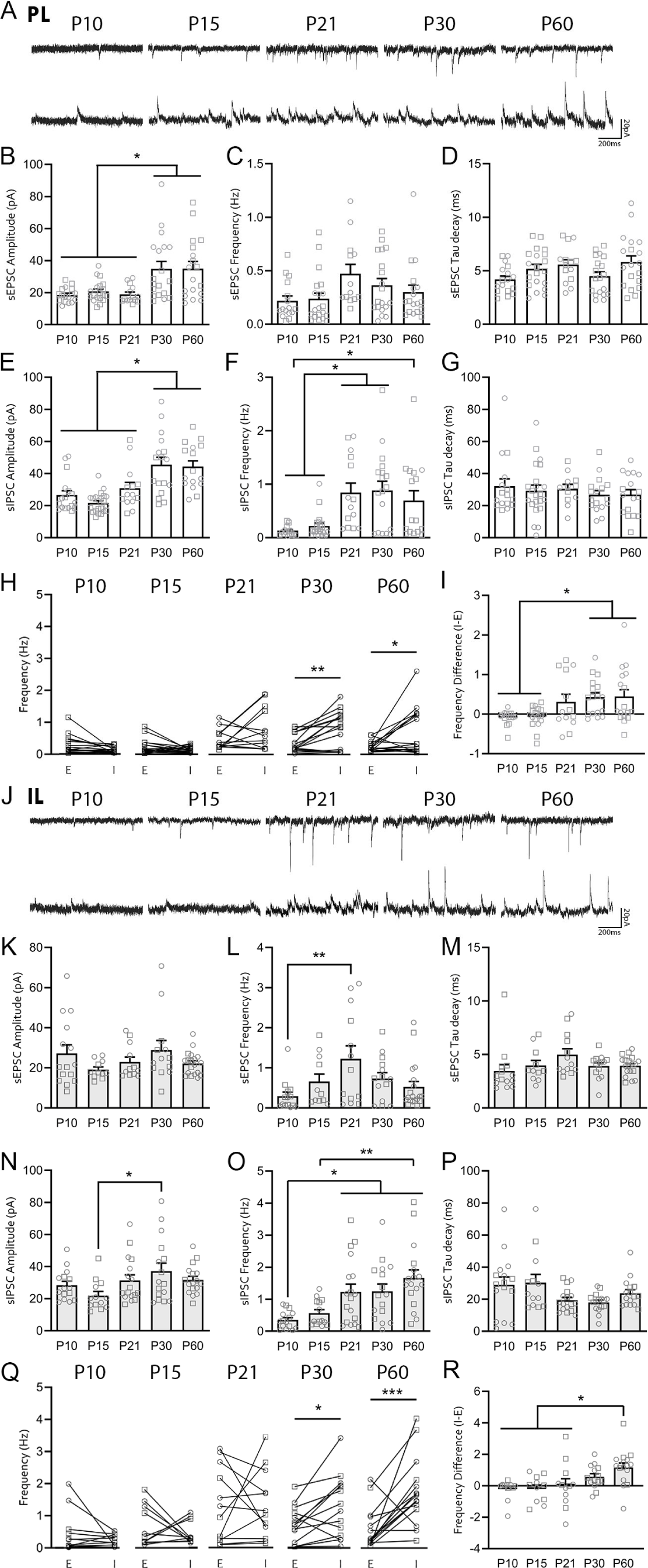
Maturation of PL and IL layer 5 Pyramidal Cell Spontaneous Activity. **A-I**. Age-dependent changes in PL layer 5 pyramidal neuron spontaneous excitatory and inhibitory postsynaptic currents (sEPSCs and sIPSCs) from P10 to P60. Example traces **(A),** sEPSC amplitude (**B**), frequency (**C**) and tau decay (**D**). sIPSC amplitude (**E**), frequency (**F**) and tau decay (**G**). **H**. Within-cell comparison of sEPSC and sIPSC frequencies. **I**. Spontaneous I-E frequency difference (calculated by subtracting sEPSC frequency from sIPSC frequency). **J-R**. Maturation of IL layer 5 Spontaneous Activity. Example traces (**J**), sEPSC amplitude (**K**), frequency (**L**) and tau decay (**M**). IL sIPSC amplitude (**N**), frequency (**O**) and tau decay (**P**). **Q**. Within-cell comparison of relative sEPSC and sIPSC frequencies. **R.** Spontaneous I-E frequency difference. Individual female datapoints are represented as circles, and males as squares. **\*** p <0.05 ** p<0.01 *** p<0.001.

Parallel analysis of the maturation of IL layer 5 spontaneous transmission showed no change in sEPSC amplitude (Fig. 4J-K) or tau decay time (Fig. 4M), but a peak in sEPSC frequency at P21 compared to P10 (Fig. 4L; one-way ANOVA p=0.017; post hoc p=0.0084). IL layer 5 pyramidal neurons showed an increase in sIPSC amplitude between P15 and P30 (Fig. 4N; one-way ANOVA p=0.048; post hoc p=0.024), and an increase in sIPSC frequency from P21 (Fig. 4O; one-way ANOVA p=0.00010; post hoc p<0.028). There were no age-dependent changes in sIPSC tau decay time (Fig. 4P). Similar to PL, we saw an increase in the relative frequency of spontaneous inhibition at P30-P60 (Fig. 4Q; paired t-test p<0.013) and increased spontaneous inhibition to excitation differences between P10-21 and P60 (Fig. 4R; one-way ANOVA p=0.0017; post hoc p<0.034), but no sex differences in this dataset (Table S1). These data indicate synchronous changes in spontaneous inhibition between PL and IL layer 5 neurons, with distinct maturation timelines for spontaneous excitation between subregions.

#### Asynchrony in the maturation of vCA1-PL and vCA1-IL presynaptic transmission

Next, we wanted to examine the maturation of pathway-specific CA1-mPFC synaptic transmission. To do that, we injected AAV1-ChR2 virus into the ventral CA1 one week prior to recording from layer 5 pyramidal neurons in the PL or IL at P15, P21, P30 and P60 (Fig. 5A). We first examined age-dependent changes in CA1-mPFC *pre*synaptic transmission. To assess glutamate release in vCA1-mPFC synapses, we analyzed EPSCs during paired pulse stimulation, whereby the relative amplitude of the second response is modulated by initial release probability. There was a significant decrease in paired pulse ratios (PPR) of vCA1-PL synapses between P21 and later ages (Fig. 5B; Mixed-effects model: main effect of age p=0.00090, interval x age interaction p<0.00010; post hoc 100ms: p<0.029; 250ms: p<0.028; Fig. 5C; 250ms: one-way ANOVA p=0.00040; post hoc p<0.0035), an effect largely driven by changes in females (two-way ANOVA p=0.036; post hoc p=0.0024). This suggests an increase in vCA1-PL release probability starting at adolescence. vCA1-IL paired pulse ratios were lower at P60 compared to P21 (Fig. 6D5D; Two-way ANOVA: interval x age interaction p=0.0036; post hoc 250ms: p=0.018; 500ms: p=0.0010; Fig. 5E; One-way ANOVA p=0.0022; post hoc p=0.020; no sex differences, Table S1), pointing to distinct presynaptic maturation trajectories for vCA1 projections to these mPFC subregions, with the vCA1-PL pathway showing an earlier profile of presynaptic changes between P21 and P30 in females.

**Figure 5.**
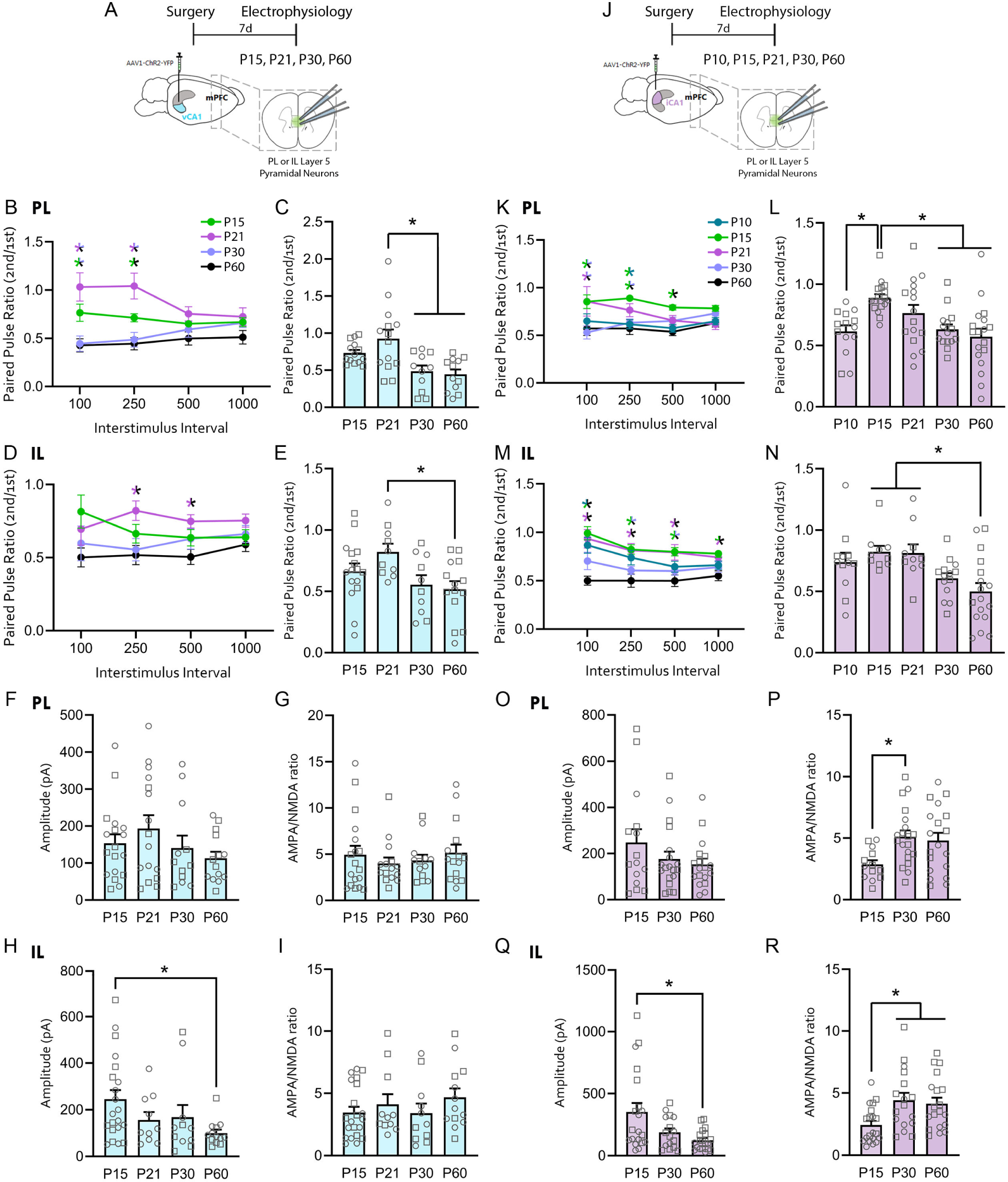
Asynchronous pre- and postsynaptic strengthening of vCA1- and iCA1-PL and IL synapses. C57BL/6J mice were injected with AAV-ChR2-YFP virus in the vCA1 (**A-I**) or the iCA1 (**J-R**) and seven days later at P10 (iCA1), P15, P21, P30 or P60 underwent patch clamp recordings from PL or IL layer 5 pyramidal neurons upon blue light LED stimulation. **A**. Experimental design for vCA1 experiments. **B-E**. Comparison of paired pulse ratio (second/first response) in PL layer 5 pyramidal neurons across intervals (**B**) and at 250ms interstimulus interval (**C**). **D-E.** Paired pulse ratios from IL layer 5 pyramidal neurons across intervals (**D**) and with 250ms interval (**E**). **F-G**. vCA1-PL layer 5 AMPAR current amplitude (**F**) and AMPA:NMDA ratios (**G**) across ages. **H-I.** vCA1-IL AMPAR amplitudes (**H**) and AMPA:NMDA ratio (**I**). **J**. Experimental design for iCA1 experiments. **K-L**. Paired pulse ratios in iCA1-PL synapses across interstimulus intervals (**K**) and at 250ms interstimulus interval (**L**). **M-N.** Paired pulse ratios in iCA1-IL synapses across intervals (**M**) and with 250ms interval (**N**). **O-R**. iCA1-PL AMPAR current amplitude (**O**) and AMPA:NMDA ratios (**P**). **Q-R**. iCA1-IL AMPAR current amplitude (**Q**) and AMPA:NMDA ratios (**R**). Individual female datapoints are represented as circles, and males as squares. **\***p <0.05. * Represents a difference between P21 and P30/P60; * between P15 and P60; * between P15 and P10; * between P10/P15 and P60; * between P15 and P30/P60; * between P15/P21 and P60; between P21 and P60, and between P15 and P21.

**Figure 6.**
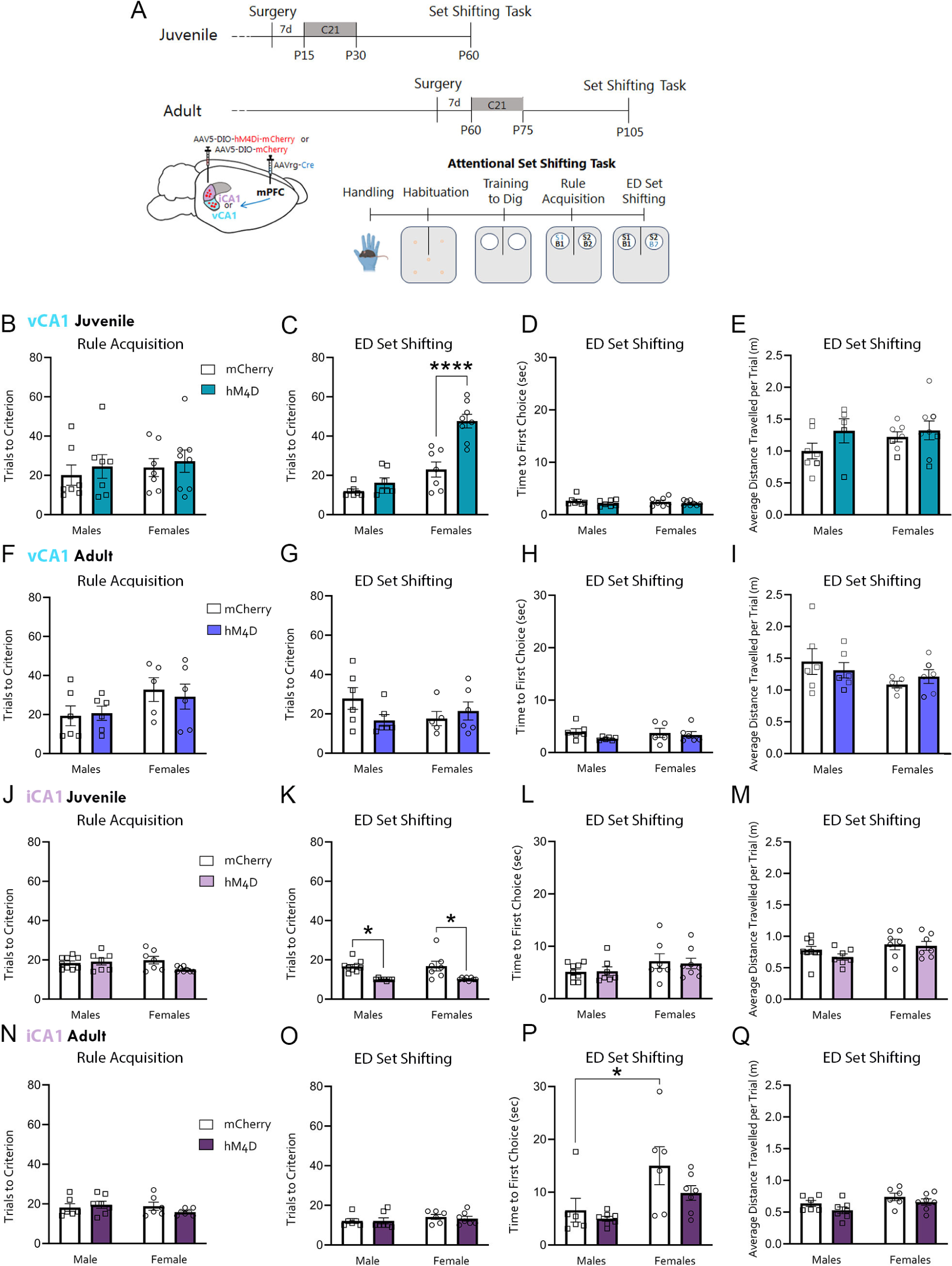
vCA1- and iCA1-mPFC inhibition during juvenility, but not adulthood, impacts extradimensional set shifting. **A.** Experimental design. Mice were injected with a retrograde AAV expressing Cre recombinase (AAVrg-Cre) in the mPFC and Cre-dependent hM4D or mCherry (AAV-DIO-hM4D-mCherry or AAV-DIO-mCherry) in either the vCA1 or iCA1 7d prior to chronic treatment with the DREADD agonist C21 for 15d either during the *juvenile* period (from P15-P30) or in *adulthood* (separate cohort receiving equivalent C21 injections from P60-P75). 30d later, animals underwent attentional set shifting^90,91^. Following handling, habituation and training to dig sessions, animals learned an association (rule acquisition) between a scent and a food reward (rewarded contingency marked in blue). The next day, animals underwent an extradimensional (ED) set shifting test in which the reward was paired with the bedding type. All groups underwent set shifting in adulthood in the absence of C21. **B-E**. vCA1 juvenile cohort. Number of trials to criterion at rule acquisition (**B**) and ED set shifting (**C**). Time to first choice (**D**) and distance travelled (**E**) during ED set shifting. **F-I**. vCA1 adult cohort. Number of trials to criterion at rule acquisition (**F**) and ED set shifting (**G**). Time to first choice (**H**) and distance travelled (**I**) during ED set shifting. **J-M**. iCA1 juvenile cohort. Number of trials to criterion at rule acquisition (**J**) and ED set shifting (**K**). Time to first choice (**L**) and distance travelled (**M**) during ED set shifting. **N-Q**. iCA1 adult cohort. Number of trials to criterion at rule acquisition (**N**) and ED set shifting (**O**). Time to first choice (**P**) and distance travelled (**Q**) during ED set shifting. Individual female datapoints are represented as circles, and males as squares. *p<0.05, ****p<0.0001.

#### Projection-specific timing signatures for vCA1- and iCA1-mPFC presynaptic maturation

We then extended this analysis to iCA1-mPFC synapses. Using an identical experimental setup, we infused the iCA1 of mice with AAV1-ChR2 virus and conducted patch clamp recordings one week later at P10, P15, P21, P30 or P60 (Fig. 5J). There was a significant peak in iCA1-PL paired pulse ratios at P15 compared to P10, P30 and P60 (Fig. 5K; Mixed-effects model: main effect of age p=0.00050; post hoc 100ms: p<0.0067; 250ms: p<0.025; 500ms: p=0.033; Fig. 5L; 250ms: one-way ANOVA p=0.00020; post hoc p<0.0079), suggesting a relative increase in presynaptic efficacy in iCA1-PL synapses from P30. When we examined iCA1-IL synapses, we saw a similar decrease in paired pulse ratio between P15-P21 and P60 (Fig. 5M; Mixed-effects model: main effect of age p<0.00010; interval x age interaction p<0.00010; post hoc 100ms: p<0.0090; 250ms: p<0.024; 500ms: p<0.030; 1000ms: p<0.012; Fig. 5N; 250ms: One-way ANOVA p=0.0015; post hoc p<0.0091). These data suggest a slight delay in the refinement of presynaptic release probability in iCA1-IL synapses (at P60) relative to iCA1-PL synapses (P30). There were no sex differences in iCA1-IL paired pulse ratios (Table S1). Together, these results reveal a differential time course for the presynaptic maturation of v/iCA1 projections to mPFC subregions, with earlier maturation of v/iCA1-PL presynaptic efficacy (P30) compared to v/iCA1-IL (P60) (see Fig. S7, left panel for a graphic summary of these data).

#### Delayed maturation of iCA1-mPFC postsynaptic efficacy

We next wanted to assess age-dependent changes in CA1-mPFC evoked *post*synaptic transmission. We recorded AMPAR currents and AMPA:NMDA ratios as a function of developmental age in vCA1- and iCA1-PL and IL synapses using the same experimental design described previously (Fig. 5A, J). We saw no change in the amplitude of AMPAR currents (Fig. 5F) or in AMPA:NMDA ratios (Fig. 5G) in vCA1-PL synapses across ages. In vCA1-IL synapses, there was a significant decrease in AMPAR response amplitude between P15 and P60 (Fig. 5H; one-way ANOVA p=0.048; post hoc p=0.033) but no change in AMPA:NMDA ratios (Fig. 5I; no sex differences, Table S1). Given the lack of changes in vCA1-mPFC postsynaptic maturation, we restricted our iCA1 analysis to P15, P30 and P60 age groups to increase experimental output. While vCA1-mPFC postsynaptic transmission remains largely stable between P15 and P60, iCA1-mPFC postsynaptic maturation undergoes marked refinement across this period. There was no change in the amplitude of AMPAR currents (Fig. 5O), but an increase in AMPA:NMDA ratios between P15 and P30 (Fig. 5P; one-way ANOVA p=0.018; post hoc p=0.019), similar to the timeline of presynaptic changes. Analysis of iCA1-IL synapses showed a decrease in AMPAR current amplitude between P15 and P60 (Fig. 5Q; one-way ANOVA p=0.0048; post hoc p=0.0044) and an increase in AMPA:NMDA ratios between P15 and P30 (Fig. 5R; one-way ANOVA p=0.0070; post hoc p<0.028), indicating synchronous maturation of postsynaptic efficacy (AMPA:NMDA ratios) of iCA1 synapses onto PL and IL subregions, which occurs after the stabilization of vCA1-mPFC postsynaptic transmission (from P15). Females showed larger iCA1-IL AMPAR current amplitude compared to males at P15 (Table S1). Overall, these data show domain-, target-, and synaptic locus-defined time signatures of synaptic maturation within the CA1-mPFC pathway. While the timing of v/iCA1 presynaptic maturation seems to be dependent on their target domain (slightly delayed maturation in IL compared to PL), vCA1 synapses display a dissociation between the timing of pre- and postsynaptic refinement (changes in PPR from P30-60 but stable AMPA:NMDA ratios across ages). Within iCA1 projections, iCA1-IL synapses show a delayed onset of presynaptic changes (P60) compared to postsynaptic changes (P30), but iCA1-PL synapses show synchronous changes in PPR and AMPA:NMDA ratios. A visual summary of the evoked CA1-mPFC synaptic maturation data is featured in Figure S7.

### Inhibition of CA1-mPFC pathways during juvenility exert opposing effects on adult cognitive flexibility

We then wanted to probe the functional implications of the CA1-mPFC maturation signatures identified by our anatomical and synaptic analyses. Critical periods in the sensory cortex are marked by structural and synaptic changes^65,67,126–130^, suggesting that the timing of CA1-mPFC synaptic changes may signal similar sensitive periods in its dependent behavior. To test this, we used an intersectional viral approach to chemogenetically inhibit either the vCA1- or iCA1-mPFC pathways at select developmental timepoints and assess its effects on a cognitive flexibility behavioral task in adulthood (Fig. 6A). We injected a retrograde AAV virus expressing Cre recombinase into the mPFC and a Cre-dependent inhibitory hM4Di AAV virus into the vCA1 or iCA1 to selectively inhibit either the vCA1- or iCA1-mPFC pathways via chronic treatment with the agonist C21, which selectively activates hM4Di receptors^131,132^ (Fig. 6A). As our electrophysiological data showed marked changes in vCA1-PL presynaptic transmission between P21 and P30 in females, we first targeted our chronic inhibition to the vCA1-mPFC pathway during the juvenile period (P15-P30) in male and female mice, and compared it to an identical manipulation during adulthood (P60-P75) (Fig. 6A). After 30d, we tested the animals in an attentional set shifting task^91^ for cognitive flexibility in the absence of C21 (Fig. 6A). In this task, mice learn to dig through two types of bedding for a food reward. During rule acquisition, the food reward is associated with a scent, regardless of bedding type (Fig. 6A). After reaching criterion, animals undergo an extradimensional set shifting test, in which the reward predictor switches to bedding type (Fig. 6A).

Mice that underwent vCA1-mPFC inhibition (vCA1 hM4D mice) during juvenility (P15-P30) showed similar rates of rule acquisition of the set shifting task performed compared to control mCherry mice in adulthood (Fig. 6B). However, vCA1 hM4D females showed an impairment in extradimensional set shifting (Fig. 6C; two-way ANOVA effect of virus and sex p<0.00010, interaction p=0.0022; post hoc p<0.00010). Importantly, juvenile-inhibited female and male vCA1 hM4D-injected animals did not differ from mCherry-injected mice in terms of time to first choice (Fig. 6D) or distance travelled (Fig. 6E) during ED set shifting, suggesting that the ED set shifting deficit seen in the vCA1 hM4D females is not driven by changes in exploration or motivation during the task. In the adult inhibition cohort, which took place between P60-P75, male and female vCA1 mCherry and hM4D mice showed similar performance at rule acquisition (Fig. 6F) and in ED set shifting (Fig. 6G-I), indicating that the ED set shifting impairment is specific to vCA1-mPFC pathway disruption during juvenility. Importantly, chronic vCA1-mPFC inhibition in either cohort did not affect performance or exploration in the elevated plus maze (Fig. S8), indicating that changes in anxiety-like behavior are not driving the ED set shifting deficit. These findings identify a sex-specific sensitive period within the vCA1-mPFC pathway during early life that is crucial for cognitive flexibility in adulthood.

To test whether the iCA1-mPFC pathway would display a similar developmental sensitivity to disruption, we conducted the same experiment but inhibiting mPFC-projecting iCA1 neurons either from P15-P30, a period in which we saw considerable iCA1-mPFC anatomical (Fig. 3I-K) and pre- and postsynaptic changes (Fig. 5K-R), or from P60-P75 (Fig. 6A), after changes had plateaued. Juvenile inhibition of the iCA1-mPFC pathway did not affect rule acquisition (Fig. 6J), but surprisingly facilitated extradimensional set shifting performance in both male and female mice (Fig. 6K; two-way ANOVA effect of virus p<0.00010; post hoc p<0.0011). iCA1 hM4D- and mCherry-injected mice showed similar time to first choice (Fig. 6L) and distance travelled (Fig. 6M) during ED set shifting. In contrast, adult iCA1-mPFC inhibition did not affect performance at rule acquisition (Fig. 6N) or extradimensional set shifting (Fig. 6O). In this cohort, we found a sex difference in the mCherry groups for latency to first choice, as females took slightly longer than males (Fig. 6P; two-way ANOVA effect of sex p=0.0046; post hoc p=0.012), but no group differences in distance travelled (Fig. 6Q). Altogether, these data highlight a convergence between pathway- (and sex-) specific synaptic and/or anatomical developmental changes and sensitivity to pathway disruption, consolidating juvenility (P15-P30) as a sensitive period for the influence of vCA1- and iCA1-mPFC pathways on adult cognitive flexibility. Importantly, they also uncovered opposite roles for juvenile inhibition of vCA1 and iCA1-mPFC pathways on adult cognitive function.

To investigate the effects of our CA1-mPFC chronic manipulations on mPFC circuit function, we next conducted whole-cell patch clamp recordings of PL layer 5 pyramidal neurons in a separate, behaviorally naïve cohort of animals (Fig. 7A). We used the same experimental design as that of the ED set shifting experiments, but recorded from PL layer 5 pyramidal cells 30d following either juvenile (P15-P30) or adult (P60-P75) inhibition of vCA1- or iCA1-mPFC pathways (Fig. 7A). Juvenile inhibition of the vCA1-mPFC pathway had no effect on PL sEPSC amplitude (Fig. 7B-C), but led to a significant decrease in PL sEPSC frequency (Fig. 7D; two-way ANOVA significant interaction p=0.0051; post hoc p=0.0050) and sIPSC amplitude (Fig. 7E; two-way ANOVA effect of virus p=0.024; post hoc p=0.0074) exclusively in females. Importantly, these changes in PL global synaptic transmission occurred in the same group that showed deficits in ED set shifting behavior (Fig. 6C). Adult inhibition of the vCA1-mPFC pathway had no effect on PL pyramidal cell spontaneous transmission (Fig. 7G-K), with the exception of an increase in sEPSC amplitude in adult vCA1 hM4D males (Fig. 7G-H; two-way ANOVA effect of virus p=0.047; post hoc p=0.018).

**Figure 7.**
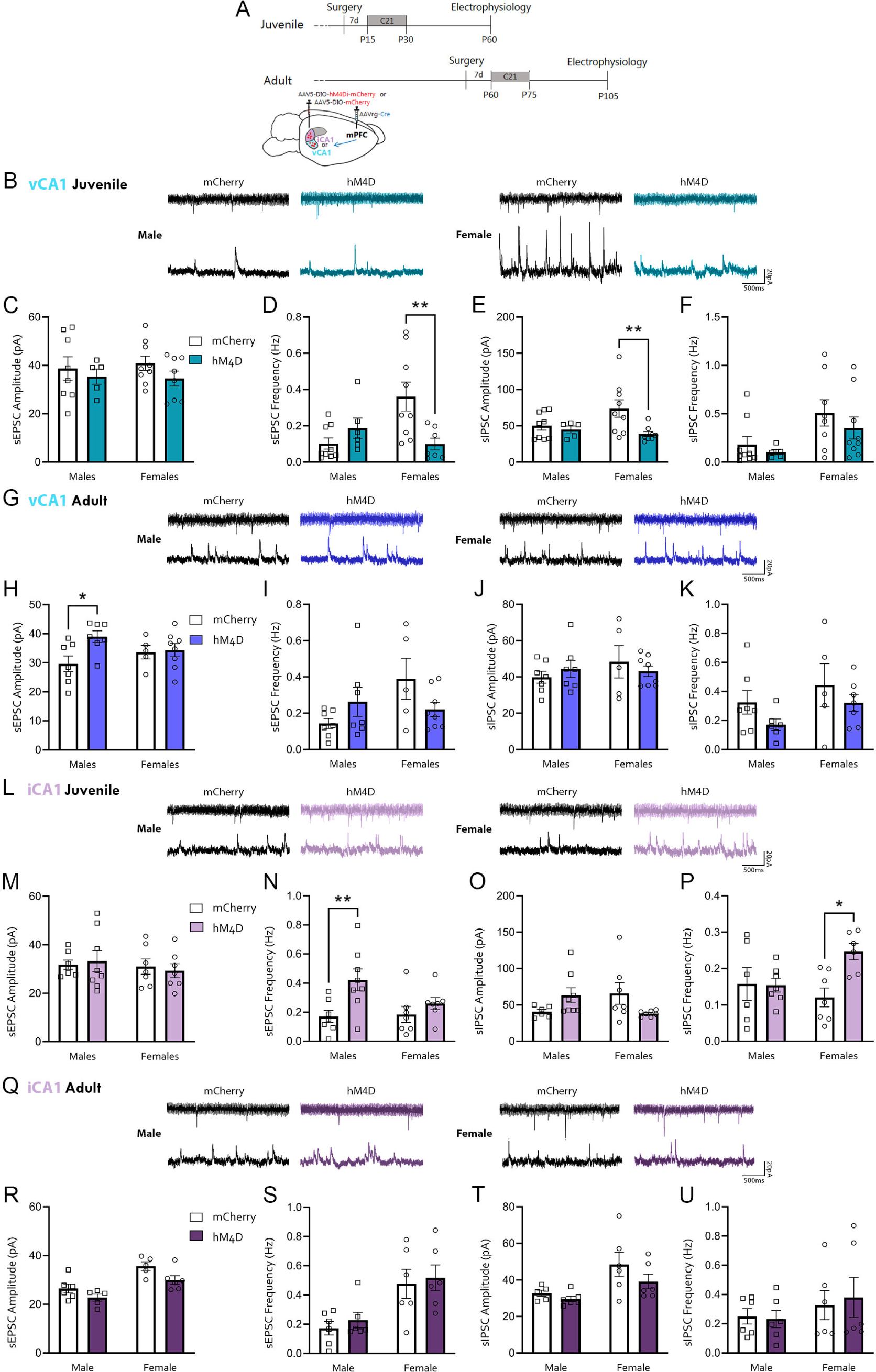
vCA1- and iCA1-mPFC juvenile inhibition impacts PL layer 5 Pyramidal Cell Spontaneous Activity. **A.** Experimental design. Mice were injected with a retrograde AAV expressing Cre recombinase (AAVrg-Cre) in the mPFC and Cre-dependent hM4D or mCherry (AAV-DIO-hM4D-mCherry or AAV-DIO-mCherry) in either the vCA1 or iCA1 7d prior to chronic treatment with the DREADD agonist C21 for 15d either during the *juvenile* period (from P15-P30) or in *adulthood* (separate cohort receiving equivalent C21 injections from P60-P75). Approximately 30d later, animals underwent patch clamp recordings from PL layer 5 pyramidal neurons in the absence of C21. **B-F**. vCA1 juvenile cohort. Example sEPSC (*top*) and sIPSC (*bottom*) traces (**B**). sEPSC amplitude (**C**) and frequency (**D**). sIPSC amplitude (**E**) and frequency (**F**). **G-K**. vCA1 adult cohort. Example traces (**G**), sEPSC amplitude (**H**) and frequency (**I**), sIPSC amplitude (**J**) and frequency (**K**). **L-P**. iCA1 juvenile cohort. Example traces (**L**), sEPSC amplitude (**M**) and frequency (**N**), sIPSC amplitude (**O**) and frequency (**P**). **Q-U**. iCA1 adult cohort. Example traces (**Q**), sEPSC amplitude (**R**) and frequency (**S**), sIPSC amplitude (**T**) and frequency (**U**). Individual female datapoints are represented as circles, and males as squares. *p<0.05, ** p<0.01.

We next examined the effects of iCA1-mPFC chronic inhibition on adult PL spontaneous transmission. Juvenile inhibition of the iCA1-mPFC pathway did not affect sEPSC amplitude (Fig. 7L-M), but led to an increase in sEPSC frequency in iCA1 hM4D males compared to mCherry controls (Fig. 7N; two-way ANOVA effect of virus p=0.0089; post hoc p=0.0085). There were no changes in sIPSC amplitude (Fig. 7O). Interestingly, iCA1-mPFC juvenile inhibition led to an increase in sIPSC frequency exclusively in females (Fig. 7P; two-way ANOVA effect of virus p=0.047, interaction p=0.037; post hoc p=0.011). Thus, the improved ED set shifting behavior seen in both sexes in the juvenile iCA1-mPFC cohort was accompanied by sex-specific changes in mPFC spontaneous transmission, with males showing changes in spontaneous excitation, and females in spontaneous inhibition. Adult inhibition of the iCA1-mPFC pathway had no effect on PL pyramidal cell spontaneous activity (Fig. 7Q-U). Overall, these data suggest that juvenile inhibition of the vCA1- and iCA1-mPFC pathways leads to disturbances in mPFC global synaptic transmission that may underlie their concurrent impact on adult cognitive flexibility.

## Discussion

Here we used a combination of viral tracing, optogenetic-assisted patch clamping, chemogenetics and behavior to examine septotemporal differences in HPC efferent projections across the lifespan, and their implications for cognitive function in adulthood. This is to our knowledge the first demonstration of a pathway-specific sensitive period for cognitive function. We found that the timing of anatomical refinement of CA1 axons varied both according to the target region and within the CA1 septotemporal axis. While CA1 projections to IL and NAc core remained stable from P10 to P60, vCA1 and iCA1 terminal density displayed distinct timing signatures in the PL, LS, NAc shell, Re, BLA, BNST and LH. Focusing on the synaptic maturation of CA1-mPFC projections, we saw an earlier onset of changes in presynaptic transmission in v/iCA1-PL compared to v/iCA1-IL synapses. Developmental changes in postsynaptic transmission also showed asynchrony among these pathways, with a marked delay within iCA1-PL and -IL synapses compared to vCA1 inputs. Finally, we uncovered a pathway- and sex-specific sensitive period for CA1-mPFC pathways during juvenility, leading to distinct modulation of cognitive flexibility in adulthood. Overall, our data consolidate a blueprint for the maturation of CA1-mPFC projections with crucial implications for behavior and the effects of circuit disturbances and/or other factors influencing circuit output from early life.

Our anatomical analysis showed unique timing of developmental signatures in vCA1 and iCA1 terminal density in the LS, NAc, BNST, LH, Re, BLA and mPFC. Notably, age-dependent refinements in these target regions were exclusive to one of the CA1-mPFC pathways, with the other pathway remaining stable for that time period. While iCA1 terminal density in LSD and LSI decreased from P21, vCA1 terminal density across these two LS subregions remained stable across ages. In rats, HPC projections shift from medial septum to LS at embryonic and perinatal (P1 and P5) stages^8,133^, but less is known about HPC-LS maturation following that period. Interestingly, in contrast to our vCA1 data, subiculum projections to the LS increase in density between P0-P30 in the rat^134^, suggesting possible species and/or subregion developmental differences.

While iCA1 terminal pruning occurs synchronously across LS subregions from P10 to later ages, projections to PL increase in density with age, similar to the LH. In rats, the first 2 to 3 postnatal weeks mark major changes in PL cytoarchitecture and an increase in dendritic arborization^135–137^, which may be informed by changes in afferent innervation, including from iCA1. While local expression of DCC/netrin guides pathway-specific anatomical innervation trajectories within mesocortical pathways^138–141^, the precise signaling molecules guiding the timing and region-specific differences in CA1 anatomical maturation still need to be identified. Furthermore, the consistency of collateral patterns seen among certain adult vCA1 subpopulations^142^ may contribute to the synchrony between vCA1 terminal changes reported here. Importantly, our functional manipulation cannot exclude the contribution of these collaterals from mPFC-projecting v/iCA1 neurons to ED set shifting performance. Future work will need to dissect the contribution of v/iCA1 subpopulations exclusively targeting the mPFC from those also projecting to amygdala^113^, LS^142^ and other regions.

Our data on age-dependent changes in spontaneous EPSCs and IPSCs at PL layer 5 showed a significant shift towards increased global presynaptic inhibition at P30. Our findings are largely consistent with work by Kroon and colleagues which described an increase in sEPSC and sIPSC frequencies between P13-16 and P26-30 in PL layer 5^136^. However, Kroon and colleagues report relatively increased excitatory responses compared to ours^136^, with differences in the age of the animals or in mouse strain/vendor possibly contributing to this disparity. Our data showed a significant shift toward PL and IL spontaneous inhibitory activity by P21-30, which is consistent with the timing of maturation of inhibitory interneurons in the mPFC between juvenility and adolescence^143–145^. Given the important role of mPFC inhibitory transmission in juvenility and adolescence to adult function^146^, dissecting the relative contributions of select input pathways to mPFC interneurons in early life will be key to elucidating the factors regulating this potential sensitive period.

We established that iCA1 and vCA1 axons arrive at the mPFC between P0 and P10 in C57BL/6J mice. This is in line with neonatal studies that report HPC oscillatory influence on the mPFC as early as P7 in rats^55,98,147^, as well as synaptic connectivity between v/iHPC and PL by P8-10^147^. Although the physiology of glutamatergic monosynaptic projections from the vCA1 to mPFC is well-known in adult mice^113,148–150^, very little was known about synaptic transmission within the CA1-mPFC circuit in early life (but see^147^). Following the initial arrival of CA1 axons, we saw marked pathway- and age-dependent changes in CA1-mPFC evoked synaptic activity. Specifically, the major changes occurring in CA1-PL transmission within the second and third postnatal weeks coincide with a P7-P13 shift from presynaptic depression to facilitation in PL layer 5 pyramidal cells following layer 2/3 stimulation in rats^151^, as well as the maturation of mPFC fast spiking interneurons and their increase in connectivity with pyramidal neurons in the second postnatal week^145^. Importantly, this period also comprises major changes in mPFC network oscillatory activity^98,145^ and strengthening of HPC-mPFC coupling between P8-10 and P16-24^46^. Moreover, changes in CA1 synaptic composition across this same timeline^152^ emphasize the need to understand the contribution of developmental changes in synapse structure to the maturation of synaptic connectivity as reported here. Interestingly, local field recordings showed that high frequency stimulation of the vHPC triggers plasticity in the mPFC in adulthood (from P65-85), but not during early (P30-40) or late (P45-55) adolescence in rats^100^, suggesting a potential further delay in the maturation of certain plasticity features within this pathway which could be reflected in our adult chronic inhibition changes in sEPSC amplitude (Fig.7H).

Our previous work had shown a distinct temporal profile for the anatomical and synaptic maturation of the mPFC-BLA pathway, which displays a delayed arrival and refinement of axons compared to CA1-mPFC projections, but synchronous evoked synaptic changes around P30^101^. Markedly, iCA1-PL, iCA1-IL and mPFC-BLA pathways display an identical timeline of postnatal maturation of AMPA:NMDA ratios, suggesting that pre- and postsynaptic maturation programs might be differentially expressed across specialized projections. The differential timeline for maturation of PL and IL projections also bears important behavioral implications given the distinct roles of these subregions in emotional and cognitive function^120–122^. Altogether, these analyses point to a high degree of specialization in the developmental profiles across brain pathways, whose guiding principles remain unknown. Given the developmental milestones taking place from juvenility to adolescence, including increased exploration and threat exposure upon leaving the nest^153^, as well as the emergence of persistent memory formation^154–159^, understanding how these time signatures relate to specialized stimulus processing and behavioral output needs further investigation.

We found that the timing of changes in CA1-mPFC synaptic transmission marked a sensitive period in the effects of pathway disruption on cognitive flexibility. For the vCA1-mPFC pathway, this effect was exclusive to females, in line with sex-specific changes in vCA1-PL PPR ratio between juvenility and adolescence, and increased vCA1-PL layer 6 terminal density in P15 females compared to males. While the origin and extent of this sex-specific effect in vCA1-mPFC maturation trajectory and sensitivity remain unclear, these data add to a growing literature showing sex differences in various aspects of HPC and mPFC maturation^160–164^, and in early life vulnerability to morphological, electrophysiological and synaptic plasticity changes in both the HPC^165–167^ and mPFC^80,168–171^. We note that the sex differences in our dataset were mostly restricted to PL maturation of spontaneous excitation and inhibition, as well as changes in vCA1-PL and iCA1-PL PPR, pointing to mPFC subregion selectivity. Nevertheless, it is important to acknowledge that given the high variability typical of developmental studies, we might have still been underpowered to detect additional sex differences in our datasets.

Juvenile inhibition of the iCA1-mPFC pathway led to an opposite effect from that of vCA1-mPFC on adult cognitive flexibility, facilitating ED set shifting performance. This suggests that vCA1- and iCA1-mPFC pathways may exert competing and opposing roles in the modulation of cognitive flexibility, a possibility that needs to be further investigated. Strikingly, juvenile iCA1-mPFC inhibition impacted adult cognitive flexibility in both males and females, whereas vCA1-mPFC juvenile inhibition only affected females. This mirrors our anatomical and electrophysiological findings, whereby vCA1-mPFC refinement was only seen in juvenile females, but was widespread across metrics and sexes for the iCA1-mPFC pathway, reinforcing a correlation between the timing of changes in pathway connectivity and sensitivity to disruption. Critically, this pattern was also reflected in the effect of juvenile chronic inhibition on mPFC spontaneous transmission, whereby juvenile vCA1-mPFC inhibition only reduced mPFC transmission in females, but mPFC spontaneous transmission was altered in both sexes following juvenile iCA1-mPFC inhibition, matching the behavioral alterations in ED set shifting performance.

Although juvenile inhibition of the iCA1-mPFC pathway led to an enhancement in ED set-shifting in both males and females, its consequences to PL spontaneous transmission were sex-specific. This adds to a growing body of work identifying sex differences in the neural basis of behavior, including in instances of equivalent behavioral output across sexes^172–175^. Additionally, pubertal onset of psychiatric disorders has been linked to changes in cognitive flexibility^176–179^, as well as changes in mPFC (and CA1^180^) perineuronal nets^181^ and increased inhibition^178^ in females, factors known to modulate sensitive periods^182–184^. Given the earlier onset of puberty in female mice relative to males^178,185^, which spans our intervention window, it will be important to uncover any potential effects of puberty on pathway-specific maturation and its modulation of cognitive flexibility.

Previous studies had implicated mPFC function in the same ED set shifting task of cognitive flexibility, with a key role for mPFC parvalbumin interneuron activity from P14-P50 in adult ED set shifting performance^146^. While this timing overlaps with that of our manipulation, the contribution of local mPFC interneurons to our findings is unknown. Furthermore, inhibition of mediodorsal thalamus (MD) from P20-P50 leads to similar ED set shifting impairments, which were accompanied by a reduction in mPFC sEPSC frequency and in the density of thalamus-mPFC projections in adulthood^91^, suggesting that both CA1- and MD-mPFC pathways exert a developmental influence on adult cognitive flexibility. Intriguingly, layer 2/3 PL pyramidal neurons receive convergent input from both MD and vHPC, while input to interneurons is biased: parvalbumin interneurons receive relatively more input from MD and VIP interneurons from the vHPC^186^. The exact interplay of these pathways and their modulation of developmental sensitivity to disruption should be further explored. Finally, a recent study showed that systemic disruption of microglia activity from P28-P37 in mice led to changes in mPFC network function and deficits in attentional set shifting rule acquisition and reversal learning in adulthood^187^. While ED set shifting was not assessed, this suggests the timing of this sensitive period might extend into adolescence, while raising the question of whether HPC input to mPFC is sensitive to microglial manipulation. Importantly, this developmental timeline is also consistent with that reported for the effects of amphetamine (P22-P31) on adult mPFC dopaminergic innervation and behavioral inhibition^141,188^, further establishing this period as one of broad sensitivity to mPFC-related circuit function and behavior.

While these studies, together with ours, support a sensitive period from juvenility to adolescence for mPFC-dependent cognitive function in adulthood, neonatal mPFC disruptions have also been linked to behavioral deficits. Transient optogenetic activation of mPFC from P7-11 leads to social, recognition and working memory deficits in juvenility and adolescence, which are accompanied by changes in mPFC morphology, electrophysiology and oscillations^189^. Additionally, serotonin transporter blockade from P2-P11 disrupts mPFC morphology, neuronal excitability and anxiety- and depression-like behavior^73^. Interestingly, the same group recently implicated serotonergic input to the mPFC in ED set shifting^190^, urging future research to explore the degree of overlap (if any) between serotonergic and v/iCA1 terminals onto mPFC pyramidal neurons. Ultimately, as research continues to refine knowledge of the topography and function of HPC projections^113,142^, the sex- and pathway-specific interrogation of the postnatal development of HPC pathways adds a crucial layer to improving our understanding of brain-behavior relationships and their contribution to the physiology of emotional and cognitive function across the lifespan.

## Methods

### Animals

C57BL/6J (Jackson Laboratory) mice were bred at the University of Toronto Scarborough and kept on a 12h light/dark cycle (lights on at 07:00 h) in a standard static cage on ¼” corncob bedding with access to food and water *ad libitum*. Date of birth was designated postnatal day (P)0, with litter sizes ranging from 2 to 11 pups. At 21 days (P21) mice were weaned and housed in same-sex littermate groups of 2 to 5 mice. Adolescence was considered at P30^191^, consistent with our previous work^101,154,192^. Littermates were randomly assigned to experimental groups, with each litter distributed across ages tested counterbalanced for sex. All experiments were conducted during the light cycle. Approximately equal numbers of females and males were used for all experiments. All animal procedures were approved by the Animal Care Committee at the University of Toronto.

### Stereotaxic surgery and viral injections

C57Bl/6J mice underwent stereotaxic surgery at P3, P8, P14, P23 or P53. 0.12-0.25µL of AAV1-CaMKIIa-hChR2(H134R)-EYFP virus was infused at the iCA1 bilaterally for each age group (P3: AP-2.5, ML-/+2.65, DV-2.5; P8: AP-2.60, ML-/+3.15, DV-3.00; P14: AP-2.65, ML-/+ 3.2, DV-3.00; P23: AP-2.7, ML-/+3.4, DV-3.00; P53: AP-2.75, ML-/+3.6, DV-3.00). For vCA1 experiments, 0.12-0.2µL of AAV1-CaMKIIa-hChR2(H134R)-EYFP virus was infused at the vCA1 bilaterally at P8, P14, P23 or P53 (P8: AP-2.46, ML-/+3.05, DV-4.60; P14: AP-1.9, ML-/+ 3.1, DV-4.6; P23: AP-2.42, ML-/+3.1, DV-4.60; P53: AP-2.53, ML-/+3.25, DV-4.5). For dCA1 experiments, 0.2uL of AAV1-CaMKIIa-hChR2(H134R)-EYFP virus was infused at the dCA1 bilaterally at P53 (AP-2.1, ML-/+1.50, DV-1.50). Hypothermia was used as an anesthetic for surgeries taken place at P3. Mice older than P8 were injected intraperitoneally (i.p.) with a combination of ketamine (100 mg/kg) and xylazine (5 mg/kg) to induce anesthesia. Once anesthetized, the head was shaved and swabbed with iodine followed by 70% ethanol. Tear gel (Alcon) was applied to the eyes to prevent dehydration. Mice were then secured in a rodent stereotaxic apparatus (Stoelting) using ear bars. Body temperature was maintained throughout surgery with a heating blanket. An incision was made down the midline of the scalp using a scalpel, the connective tissue was excised, and then the skull was cleaned with sterile phosphate buffered saline (PBS, pH=7.4). An autoclaved cotton-tip applicator was briefly submerged in 30% hydrogen peroxide and gently applied to the skull surface to identify bregma. After surgery, mice recovered for one week and then were transcardially perfused or underwent slice electrophysiology (details described below). For the retrograde tracing surgeries, mice at P0, P5 or aged 8-9 weeks underwent stereotaxic surgery (as above) infusing, bilaterally, red Lumafluor RetroBeads™ at the mPFC (P0: AP+0.3, ML-/+0.2, DV-0.75; P5: AP+1.07, ML-/+0.25, DV-2.6; Adult: AP+2.0, ML-/+0.3, DV-2.8). Five days after surgery, mice underwent transcardial perfusion (details described below). For the set shifting experiments, mice at P8 or P53 underwent stereotaxic surgery (as described above) infusing bilaterally, pAAV5-hSyn-DIO-hM4D(Gi)-mCherry or pAAV5-hSyn-DIO-mCherry at the vCA1or iCA1 (coordinates for P8 and P53 above) and pENN-AAV-hSyn-Cre-WPRE-hGH (Retrograde) at the mPFC (P8: AP+1.52, ML-/+0.26, DV-2.5; P53: AP+2.0, ML-/+0.3, DV-2.8). After surgery, mice recovered for one week prior to commencing chronic C21 injections (procedure described below). We chose a chemogenetic intersectional approach in these experiments because of challenges related to optic fiber implantation in very young mice (in the juvenile group), in addition to limitations of chronic light stimulation in optogenetic experiments.

### Perfusions and sectioning

Mice were injected with Avertin (250mg/kg, i.p.) and once under deep anesthesia, transcardially perfused with approximately 50mL of room temperature 0.1M PBS (pH=7.4), followed by approximately 80mL of cold 4% paraformaldehyde (PFA). Brains were extracted and stored overnight at 4°C in 4% PFA. Brains were subsequently sectioned at 50µm in the coronal plane (VT1000, Leica) and stored at -20 °C in a cryoprotectant solution comprised of 60% glycerol and 0.01% sodium azide in PBS.

### Immunohistochemistry

Immunostaining was performed on free floating sections. Sections were washed in 0.1M PBS (3 x 5 min each) and then incubated in blocking solution comprising 5% normal goat serum and 0.25% Triton X-100 in 0.1M PBS for 30 min. For the amplified dCA1, vCA1 and iCA1 brains, amplification of the viral signal was achieved by incubating sections with polyclonal chicken anti-GFP (1:1000, #ab13970; Abcam, RRID: AB_300798) primary antibody diluted in blocking solution. Sections were incubated overnight at 4 °C on a rotary shaker under gentle agitation. On the following morning, sections were incubated in goat anti-chicken Alexa 488 (1:1000, #A11039, Invitrogen, RRID: AB_2534096) secondary antibody for 2 hours and Hoechst 33342 nuclear dye (1:1000 diluted in 0.1M PBS; #H1399; ThermoFisher). Sections were then rinsed in 0.1M PBS, mounted onto gelatin-coated slides and coverslipped with Permafluor anti-fade mounting medium (#TA030FM; Thermo Scientific).

### Anatomical maturation analysis

Brains were sectioned (50µm thickness) with a VT1000S vibratome (Leica). All sections were stained with nuclear dye Hoechst 33342 (1:1000) for 10 min at room temperature. Sections were mounted onto slides, air dried, and coverslipped with anti-fade mounting medium (as described above) and imaged using a Nikon Eclipse Ni-U Upright epifluorescent microscope. iCA1 and vCA1 projection-associated EYFP fluorescence was quantified by measuring average pixel intensity for each region of interest and subtracting the background/basal measure of intensity. For the vCA1/iCA1 measurements (Figure 1), high fluorescence intensity at the injection site prevented quantification of individual neurons. The terminal density analyses (Figures 2, 3, S1) exclusively captured axon fluorescence, as we did not see cell bodies in any of the terminal target images. Internal baseline measures of intensity for each section were taken from the motor cortex/frontal associative cortex which are devoid of temporal hippocampal projections^114^. An average intensity difference was calculated from both hemispheres using ImageJ/FIJI software (NIH, MD, USA; https://imagej.net/software/fiji/). Regions of interest included: the prelimbic cortex (PL) (layer 2/3, 5 and 6), infralimbic cortex (IL) (layer 2/3, 5 and 6), nucleus accumbens (NAc) (shell and core), lateral septum (LS) (dorsal, ventral and intermediate nucleus), basolateral amygdala (BLA), lateral hypothalamus (LH), thalamic nucleus reuniens (Re) and the bed nucleus of the stria terminalis (BNST). Whole area intensity was measured for each region. Comparisons were made of vCA1/iCA1 projection-associated EYFP intensity at these regions among the different age groups. All calculated intensities were normalized with EYFP intensity in vCA1 or iCA1 injection site.

To ensure viral spread is restricted only to, and covers the whole of the vCA1 or iCA1, the Cavalieri method [(V_c_ = *d*(Ʃ(y_i_)), where V_c_ = estimated volume (mm^3^), *d* = distance between sections (µm) and y = area of EYFP infection or vCA1/iCA1 per section (mm^2^)] was used to estimate, for each age, viral spread and total vCA1/iCA1 volume. The iCA1 is defined as the dorsal CA1 (dCA1^i^) region (pyramidal, oriens and radiatum layers) that begins from bregma -2.70mm and terminates at -3.88mm labelled in Franklin & Paxinos mouse atlas^193^. For sections that do not have the CA2 region that easily delineates the dCA1^i^ on the dorsal-ventral axis within bregma -2.70 to -3.88mm, the dCA1^i^ consisted of the CA1 area above the rhinal fissure/bottom limit of the perirhinal cortex^193^. This boundary follows the lower border of the intermediate CA1 set by Dong et al. (2009) and Fanselow & Dong (2010). Since these authors found that their iCA1 region and temporal/posterior dCA1 region follow the same pattern of gene expression^22^, for this study, the iCA1 encompasses both Dong et al. (2009)’s posterior dCA1 and iCA1. Estimated volume of EYFP and vCA1/iCA1 were compared between each age group to corroborate the reduction of hippocampal mass and the restriction of viral spread at earlier ages^101^. Results are expressed as percentage of estimated EYFP volume and vCA1/iCA1 volume.

For the quantification of cells infused with Lumafluor RetroBeads™ cells (Figure S3), images were obtained a Nikon Eclipse Ni-U Upright epifluorescent microscope and manually counted using ImageJ/FIJI software (NIH, MD, USA; https://imagej.net/software/fiji/).

### Slice electrophysiology

Mice were anesthetized with isoflurane and perfused transcardially with ice-cold (0°C– 4°C) sucrose solution composed of (in mM): 197 sucrose, 11 glucose, 2.5 KCl, 1.25 NaH2PO4, 25 NaHCO3, 0.4 ascorbate, 1 CaCl2, 4 MgCl2 and 2 Na pyruvate. Acute 350 μm coronal slices of PFC were obtained from dissected brains using a VT1000S vibratome (Leica) and then incubated at 35°C for 25 min in the same solution, but with 50% standard artificial cerebral spinal fluid (ACSF) composed of (in mM): 120 NaCl, 2.5 KCl, 1.25 NaH2PO4, 25 NaHCO3, 11 glucose, 2 CaCl2, 1 MgCl2 and 2 Na pyruvate. Following recovery, slices were maintained at room temperature in standard ACSF. Whole-cell voltage-clamp recordings were obtained from principal pyramidal neurons in layer 5 PL or IL using borosilicate glass electrodes (3–5 MΩ). Electrode internal solution was composed of (in mM) of the following: 130 Cs-methanesulfonate, 10 HEPES, 0.5 EGTA, 8 NaCl, 4 Mg-ATP, 1 QX-314, 10 Na-phosphocreatine, and 0.4 Na-GTP. Pyramidal excitatory neurons in the mPFC were discriminated on the basis of soma size, high capacitance (>200 pF), and slow EPSC kinetics, under the rationale that no interneuron subtypes possess all three of these characteristics. CA1 axon terminals were stimulated using illumination with a digital multimirror device (Polygon400, Mightex) and a TTL-pulsed microscope objective-coupled light-emitting diode (LED, 470 nm, 14 mW/mm2, Mightex). This pulse intensity evoked maximal response amplitudes at all ages. For analysis of AMPA/NMDA ratios, we applied 1 μM TTX (#HB1035, HelloBio) and 100 μM 4-AP (#AB120122, Abcam) for more stringent isolation of monosynaptic transmission ^101,194,195^. Paired pulse evoked EPSC and spontaneous EPSCs and IPSCs were conducted in standard ACSF described above. Paired pulse evoked EPSC were recorded using a low chloride internal (TEA-Cl 5mM). Data were low-pass filtered at 3 kHz (evoked) and 10 kHz (spontaneous) and acquired at 10 kHz using Multiclamp 700B and pClamp 10 (Molecular Devices). All stimulation was conducted at 0.1 Hz to avoid inducing synaptic plasticity. Series and membrane resistance was continuously monitored, and recordings were discarded when these measurements changed by >20%. Recordings in which series resistance exceeded 25 MΩ were rejected. Detection and analysis of sEPSCs and sIPSCs were performed blind to experimental group using MiniAnalysis (Synaptosoft). EPSC decay time constants (τ decay) were obtained by standard exponential curve fitting of averaged current traces in Clampfit (Molecular Devices). AMPA/NMDA ratio was calculated as the peak EPSC at - 70 mV divided by the EPSC amplitude at 100ms after current onset at +40 mV^101,194^.

### C21 injections

Animals were injected for 16 days consecutively with DREADD agonist compound 21 (C21; 2mg/kg, 0.5mg/mL, i.p.; #HB6124; HelloBio)^131^, twice a day^91^, once at 8:30 am ± 1 hour and once at 4:30 pm ± 1 hour, ensuring an 8-hour gap in between. The juvenile group was injected from P15-P30, and the adult group was injected from P60-P75. These cohorts underwent either attentional set shifting or slice electrophysiology (whole cell patch clamp recordings of sEPSCs and sIPSCs as described above) 30d later in the absence of C21.

### Behavioral testing

#### Attentional Set Shifting Task

##### Apparatus

A rectangular chamber made of white plexiglass (14.5 cm x 16.5 cm) was separated by a 20 cm long x 16.5 cm wide x 0.5 cm thick plexiglass wall with two 19.2 cm wide x 17.8 cm long adjoining sliding doors on either lateral side to enclose the mouse in the left or right upper quadrant. The apparatus was elevated 41 cm off the floor with an Imaging Source DMK 22AUC03 1.1.3 IR camera mounted 88.4 cm directly above it using a wall mount rack. An external LED lamp (280 Lux) was positioned adjacent to the camera and 87.4 cm above the chamber to evenly distribute light. Two round ceramic saucers each measuring 8.9 cm in diameter were placed in the chamber at each of the quadrants during testing (at a 2.8 cm distance from each lateral wall and 7.6 cm away from the sliding door). The saucers were filled with one of two types of bedding [counterbalanced; bark (Zoo Med Repti Bark) or cellulose paper (Biofresh 1/8” Pelleted Cellulose) bedding], mixed with either a garlic or cinnamon powder (McCormick), one of which predicted the reward location. Four different variations of scent–bedding mixtures were used: bark–cinnamon, bark–garlic, paper–cinnamon and paper–garlic.

##### Procedure

We used a variation of the protocol described by the Kellendonk group^90,91^. Animals were assessed in adulthood at P56-P61 for the juvenile group and at P101-P106 for the adult group. Animals started undergoing food deprivation on the first day of handling (Day 1). Mice were handled for 2 days (twice per day for 2 minutes, Days 1-2). On Day 2, after the second handling session, mice underwent one 10-minute habituation session in which they were exposed to the experimental apparatus in the presence of cheerios spread across the chamber to encourage exploration (no saucers present). Starting on the day after habituation (Day 3), mice underwent two days of training to dig (6 trials/day), when they learn to dig in the saucer filled with bed-o-cob bedding for a cheerio reward. The bedding amount was increased incrementally over the six daily sessions until the cheerio was fully covered. On the following day (Day 5), animals underwent rule acquisition. During rule acquisition, animals were exposed to two saucers displaying a pseudorandom, counterbalanced combination of one of the two beddings and scents, one of which contained a cheerio reward. The reward was paired with one of the scents (rule; counterbalanced) for the first five warm-up trials until they found the cheerio. From trial six onwards, once the animal’s snout crossed the edge of the saucer, this was considered a decision, which was followed by enclosing the animal within the side of the chamber for 30 seconds using a sliding door. Trials ran until animals reached the criterion of 8 correct choices within 10 consecutive trials, taken as a proxy for learning the scent–reward association. The next day (Day 6), mice underwent extradimensional (ED) set shifting. During ED set shifting, a similar procedure from rule acquisition was followed, but the bedding medium (instead of the scent) predicted the cheerio location. Trials ran until animals reached the criterion of 8 correct choices within 10 consecutive trials. Animals that did not meet this criterion before reaching 70 trials were excluded from the experiment. Scent and bedding types were counterbalanced between animals. Trials were recorded using the Imaging Source DMK 22AUC03 1.1.3 IR camera and distance travelled and time to first choice were quantified using ANYmaze software (Stoelting, USA). Trials to criterion were quantified manually by a blind experimenter and validated through ANYmaze. Two juvenile hM4D animals showed an outlier profile for distance travelled during set shifting (3.78m and 3.98m), which was solely driven by exploration after choice (while eating the food reward) and were therefore excluded from that analysis.

### Quantification and statistical analysis

Data are presented as mean ± SEM with n shown as the number of animals for histological experiments, or number of cells followed by number of animals in parenthesis for electrophysiological experiments. Due to space limits, only significant p is reported in the main text, with all other detailed statistical analyses (including sex differences and nested ANOVAS) reported in Table S1. All statistical analyses were performed in GraphPad Prism® version 9 or Jamovi version 2.7.6 software through one-way ANOVA, two-way repeated-measures or nested ANOVAs, where appropriate. Omnibus test results (F values) are reported in Table S1. To test for group differences following ANOVA, we applied the software default of Tukey’s, Bonferroni’s or Šídák’s-corrected post hoc tests for all pairwise comparisons following non-repeated and repeated measures ANOVA, respectively, and significant results were denoted by asterisks in associated figures. Potential sex differences were first assessed using a two-way, repeated measures ANOVA and in the absence of effects, data were collapsed across these variables for subsequent analyses. Outliers were detected using the software outlier test (Grubbs’ Test or the extreme studentized deviant method) and removed from the dataset prior to running group and/or post hoc comparisons.

## KEY RESOURCES TABLE

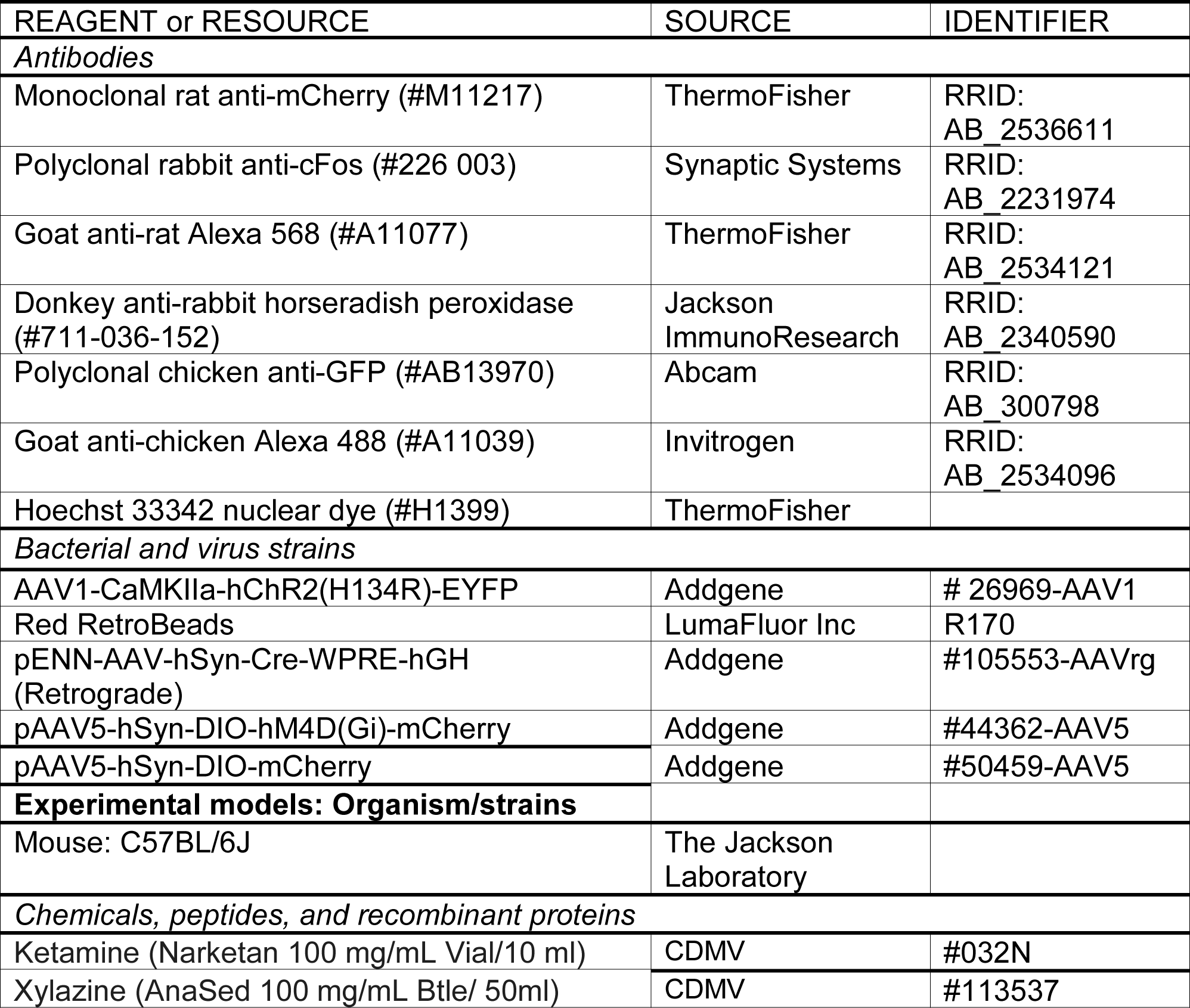
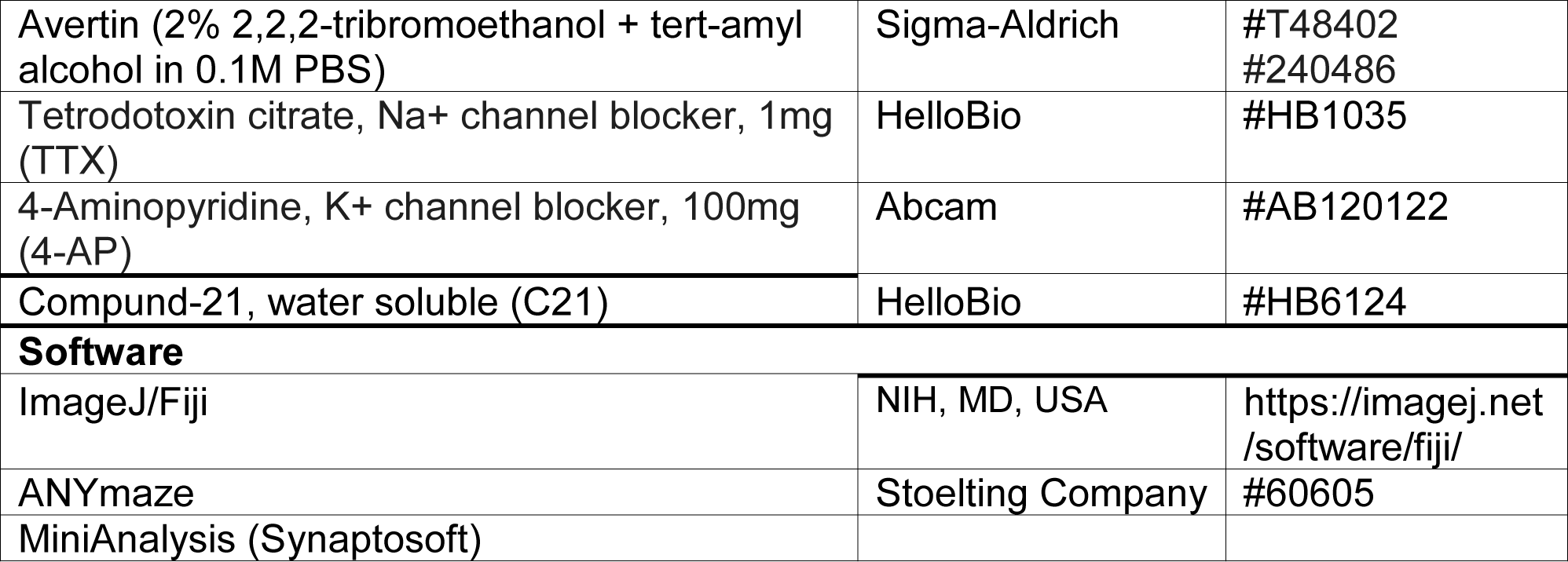

## Acknowledgements

This work was supported by a Natural Science and Engineering Council of Canada (NSERC) CGS-D (CGSD3 - 534884 – 2019) for AC-S, a grant from the NIH (R01MH126105) to CA, as well as grants from the Natural Science and Engineering Council of Canada (RGPIN-2017-06344), SickKids Foundation and Canadian Institutes of Health Research (CIHR) – Institute of Human Development, Child and Youth Health (NI19-1132R), CIHR (PJT 399790), and Human Frontier Science Program Organization (CDA00009/2018 and RGY0072/2019) to MA-C. We acknowledge resources and support from the Centre for the Neurobiology of Stress (CNS) Core Facility at the University of Toronto Scarborough, and thank Durga Acharya and Bruno Chue for their help in the facility. A Canada Foundation for Innovation grant (#493864) was used to establish the Centre for the Neurobiology of Stress (CNS) Core Facility.

## Supplemental Figures and Legends

**Figure S1.**
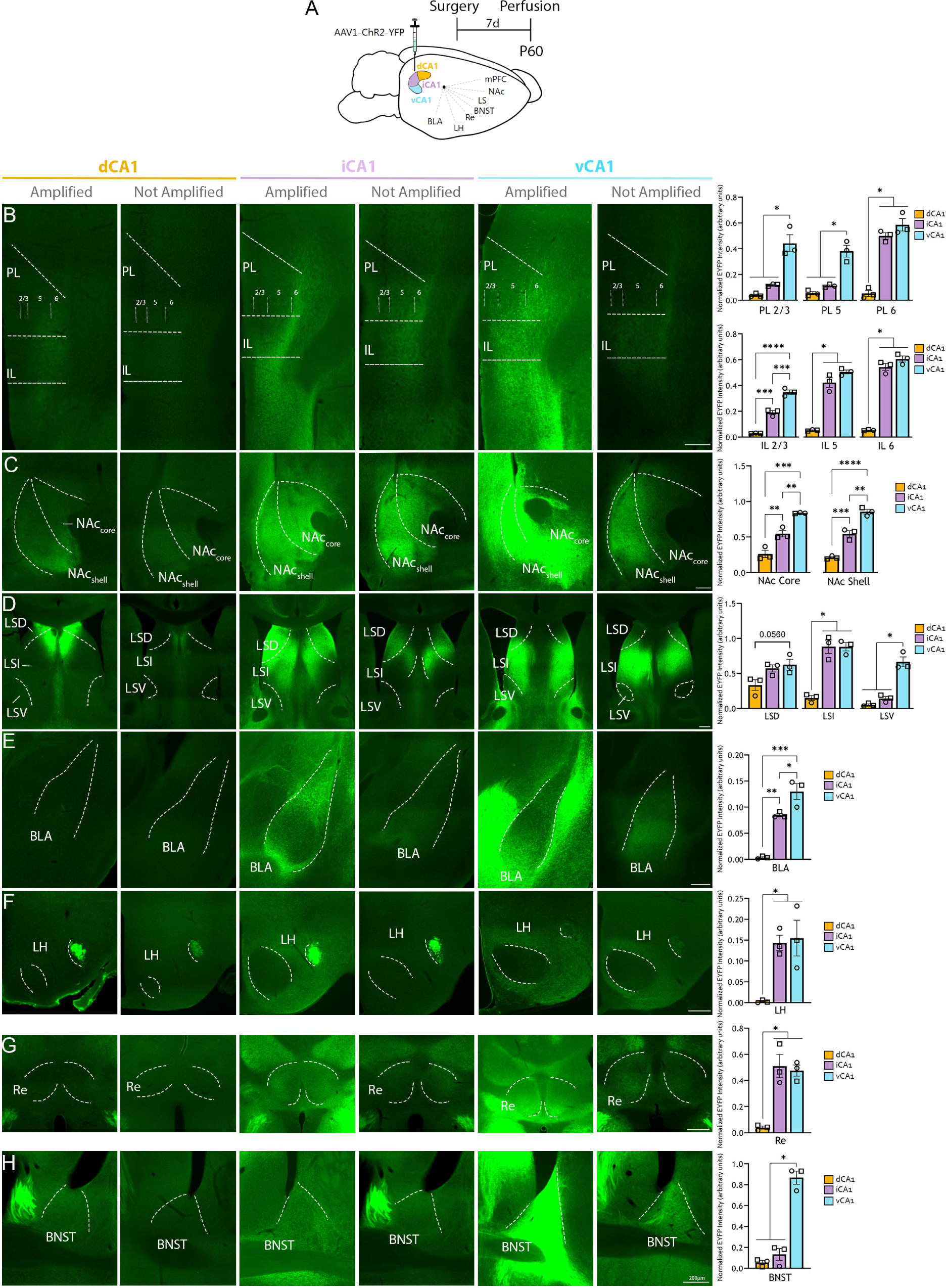
Comparison of projection patterns from dorsal, intermediate and ventral CA1 in adult mice. **A**. C57BL/6J mice were injected with AAV-ChR2-YFP virus in the dorsal, intermediate or ventral CA1 and perfused seven days later at P60 for tissue and image processing. **B-H.** Example images and quantification of differences between projections from dCA1 (left image columns) and those from intermediate iCA1 (middle image columns) and vCA1 (right most columns), with the iCA1 and vCA1 projections showing largely similar patterns across examined regions. The left most column of each CA1 set of images (dCA1 amplified, iCA1 amplified, vCA1 amplified) underwent immunohistochemistry against GFP to amplify viral signal. All other (not amplified) images show direct fluorescence under identical settings. Graphs on the right show semi-quantification of terminal density of amplified images for each of the three pathways across all regions. BNST = bed nucleus of the stria terminalis, BLA = basolateral amygdala, IL = infralimbic cortex, LH = lateral hypothalamus, LSD = dorsal lateral septum, LSI = intermediate lateral septum, LSV = ventral lateral septum; NAc = nucleus accumbens, PL= prelimbic cortex, Re = thalamic nucleus reuniens. Scale bar: 200μm. n=3 per projection. Individual female datapoints represented as circles, male as squares. *p<0.05, **p<0.01, ***p<0.001, ****p<0.0001.

**Figure S2.**
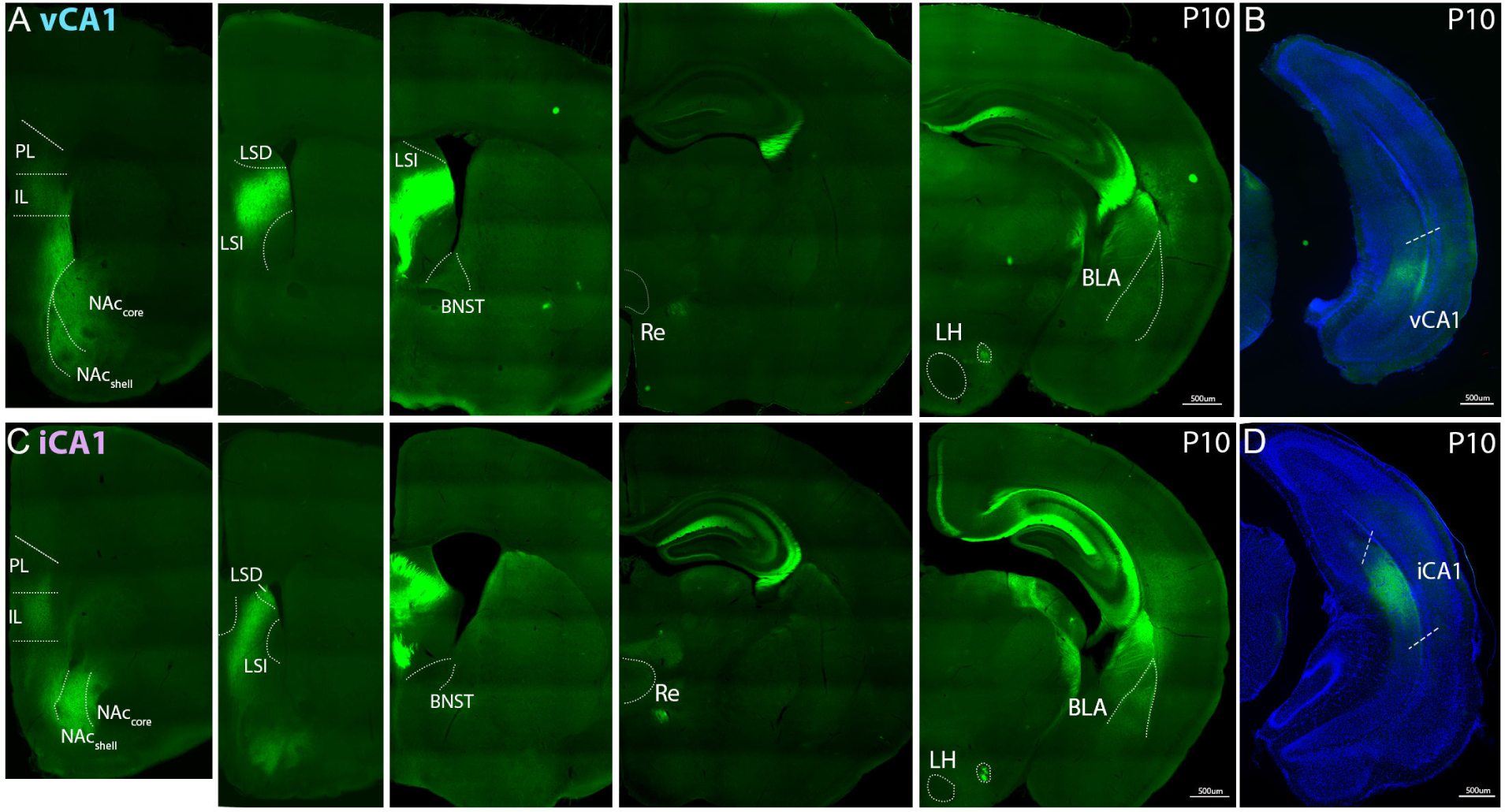
iCA1 and vCA1 axons are already present in all target areas by P10. C57BL/6J mice were injected with AAV-ChR2-YFP virus in the ventral (vCA1; **A-B**) or intermediate (iCA1; **C-D**) CA1 at P3 and perfused seven days later at P10 for tissue and image processing. **A** and **C** show terminal projections for each pathway (**A**: vCA1; **C**: iCA1), whereas **B** and **D** show viral expression at the respective infusion site. vCA1 and iCA1 axons were already visible at P10 in the prelimbic cortex (PL), infralimbic cortex (IL), nucleus accumbens (NAc) core and shell, dorsal lateral septum (LSD) and intermediate lateral septum (LSI), thalamic nucleus reuniens (Re), basolateral amygdala (BLA), bed nucleus of the stria terminalis (BNST), and lateral hypothalamus (LH). Scale bar: 500μm.

**Figure S3.**
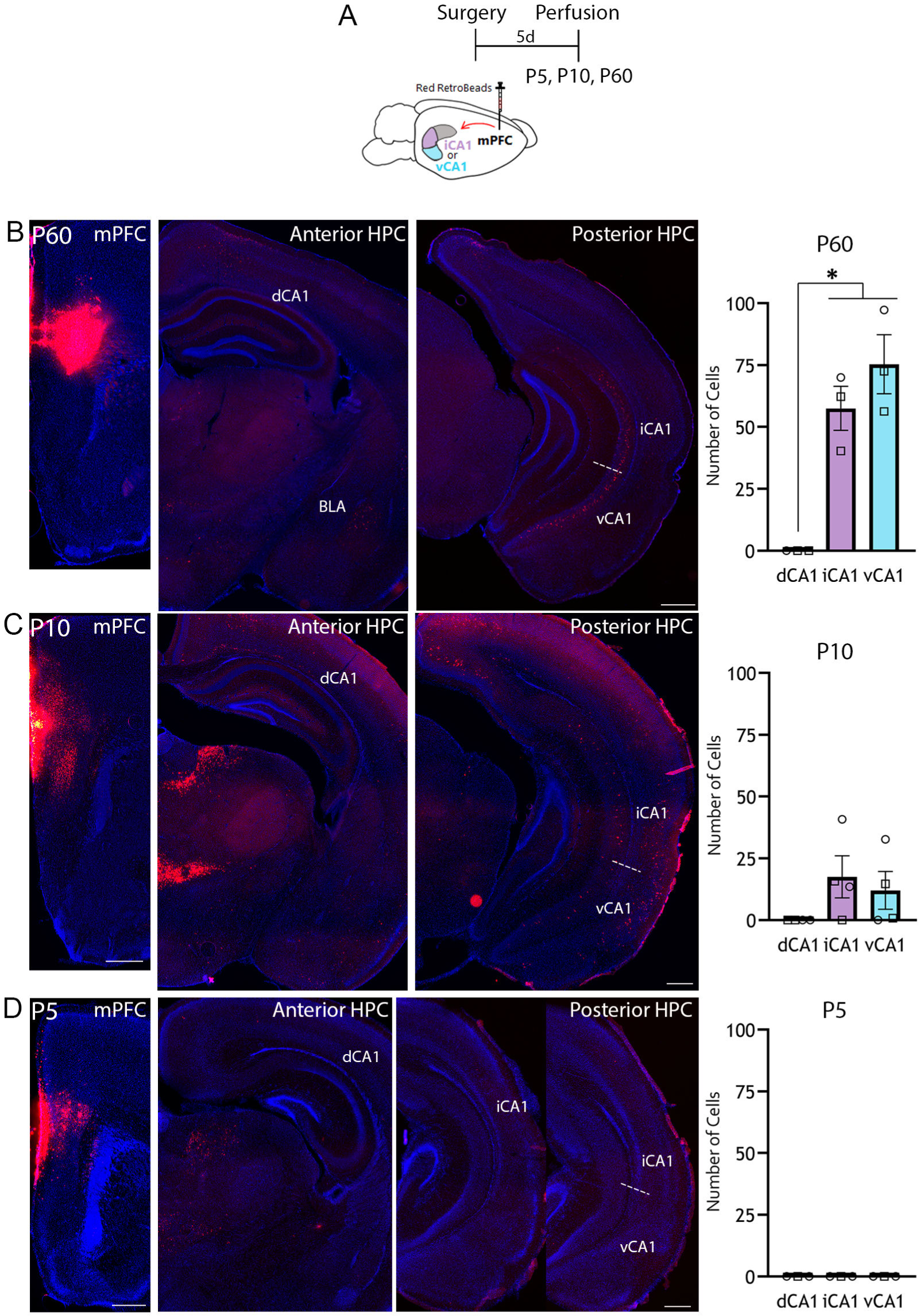
CA1 terminals arrive at the mPFC between P0 and P10 in C57BL/6J mice. **A**. Schematic of experiment. C57BL/6J mice were infused with red Lumafluor RetroBeads™ unilaterally at the medial prefrontal cortex (mPFC). Mice were transcardially perfused 5 days following surgery at P5, P10 or P60. **B**. Sample image of the mPFC infusion site in an adult P60 brain (*left most image*) shows absence of red soma signal at the dorsal CA1 (dCA1) but labeled cells in the basolateral amygdala (BLA) (*center image*) and in the intermediate CA1 (iCA1) and ventral CA1 (vCA1) (*right most image*). **C**. Sample image of the mPFC infusion site in a P10 brain (*left most image*) showing absence of red soma signal at the dCA1 but labeled cells in the BLA (*center image*), as well as in the iCA1 and vCA1 (*right most image*). **D**. Sample image of the mPFC infusion site in a P5 brain (*left most image*) showing absence of red soma signal at the dCA1, iCA1 or vCA1 (*right most images*). Graphs on the right show quantification of the number of labeled cells in the dCA1, iCA1 and vCA1. n=3 for each age. Individual female datapoints represented as circles, male as squares. *p<0.05. Scale bar: 500μm.

**Figure S4.**
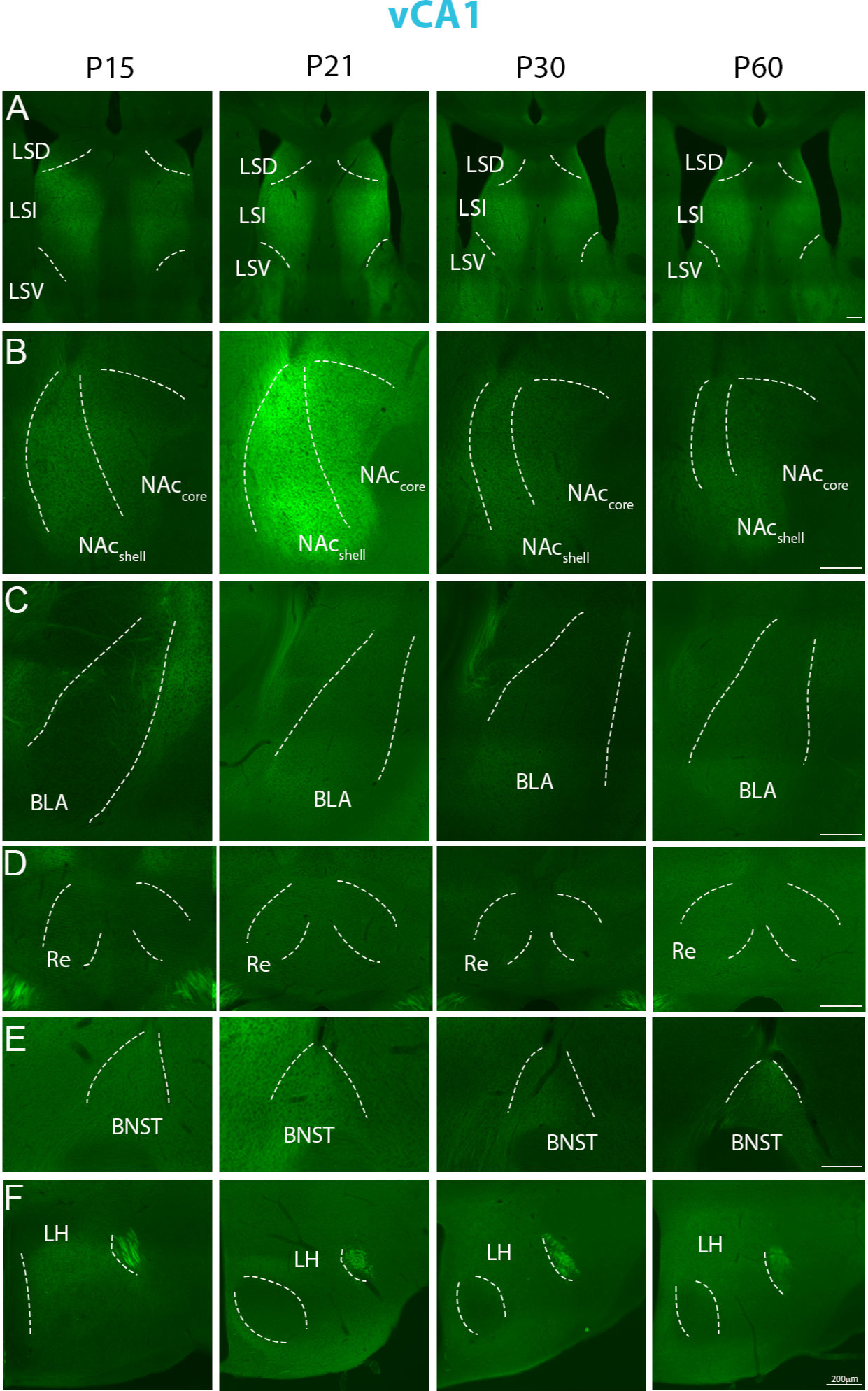
Example images showing developmental changes in vCA1 terminal density. C57BL/6J mice were injected with AAV-ChR2-YFP virus in the vCA1 and perfused seven days later at P15, P21, P30 or P60 for tissue and image processing. Images show vCA1 terminal expression across ages in the dorsal lateral septum (LSD; **A**), ventral lateral septum (LSV; **A**), intermediate lateral septum (LSI; **A**), nucleus accumbens (NAc) core (**B**) and shell (**B**); basolateral amygdala (BLA; **C**), thalamic nucleus reuniens (Re; **D**), bed nucleus of the stria terminalis (BNST; **E**), and lateral hypothalamus (LH; **F**).

**Figure S5.**
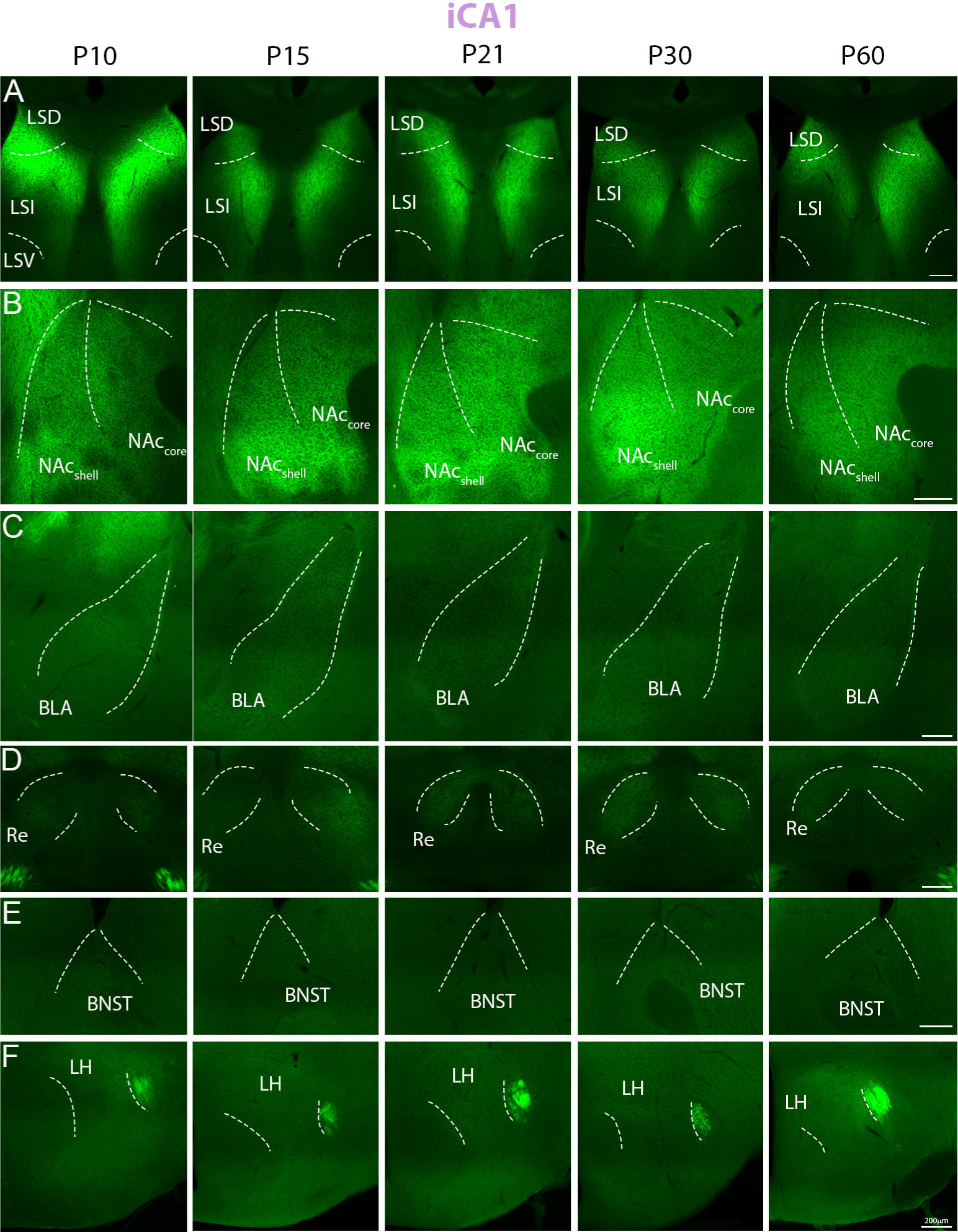
Example images showing developmental changes in iCA1 terminal density. C57BL/6J mice were injected with AAV-ChR2-YFP virus in the iCA1 and perfused seven days later at P10, P15, P21, P30 or P60 for tissue and image processing. Images show iCA1 terminal expression across ages in the dorsal lateral septum (LSD; **A**), intermediate lateral septum (LSI; **A**), nucleus accumbens (NAc) core (**B**) and shell (**B**); basolateral amygdala (BLA; **C**), thalamic nucleus reuniens (Re; **D**), bed nucleus of the stria terminalis (BNST; **E**), and lateral hypothalamus (LH; **F**). LSV = ventral lateral septum.

**Figure S6.**
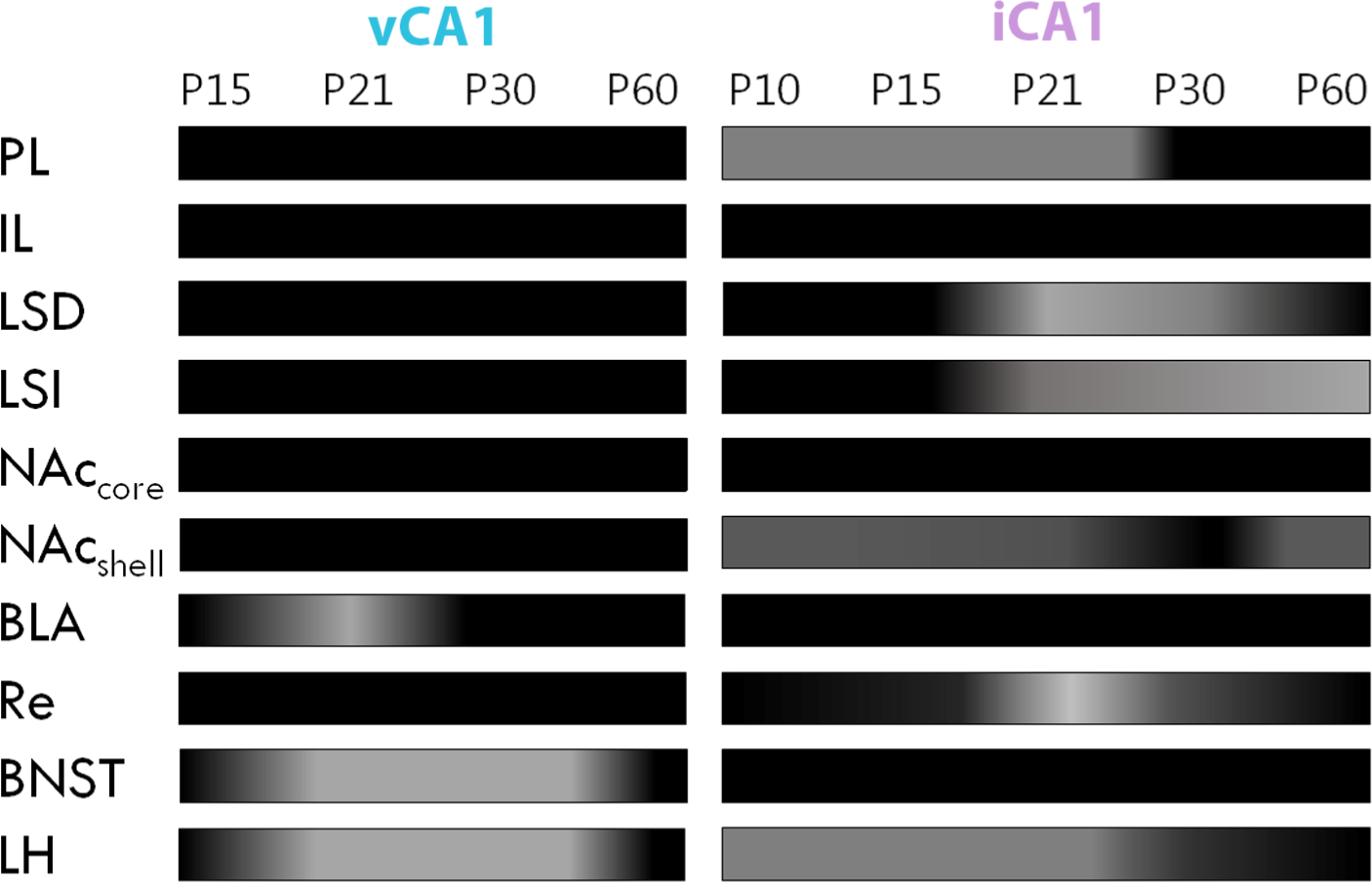
Graphic representation of anatomical maturation data for vCA1 and iCA1 projections. Rectangular heatmaps represent relative changes in terminal density across respective ages, whereby darker color represents an increase. PL = prelimbic cortex, IL = infralimbic cortex, NAc = nucleus accumbens, LSD = dorsal lateral septum, LSI = intermediate lateral septum, Re = thalamic nucleus reuniens, BLA = basolateral amygdala, BNST = bed nucleus of the stria terminalis, LH = lateral hypothalamus.

**Figure S7.**
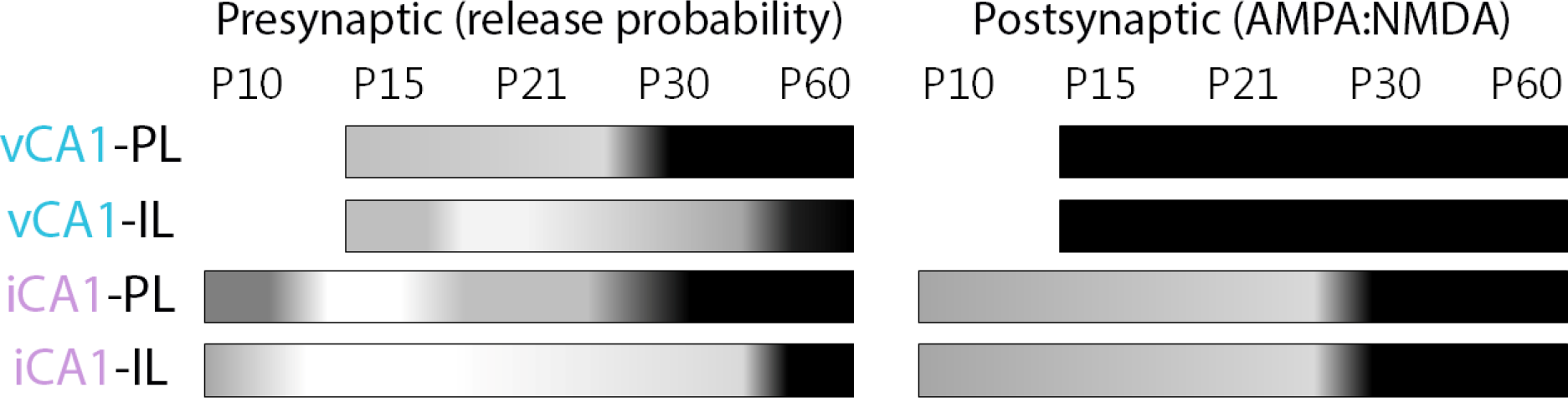
Graphic representation of evoked synaptic maturation data for vCA1- and iCA1- PL and IL pathways. Rectangular heatmaps represent relative changes in presynaptic (**Left**; or release probability, the inverted function of the paired pulse ratios) and postsynaptic efficacy (**Right**, represented as AMPA:NMDA ratios) across respective ages, whereby darker color represents an increase. PL = prelimbic cortex, IL = infralimbic cortex.

**Figure S8.**
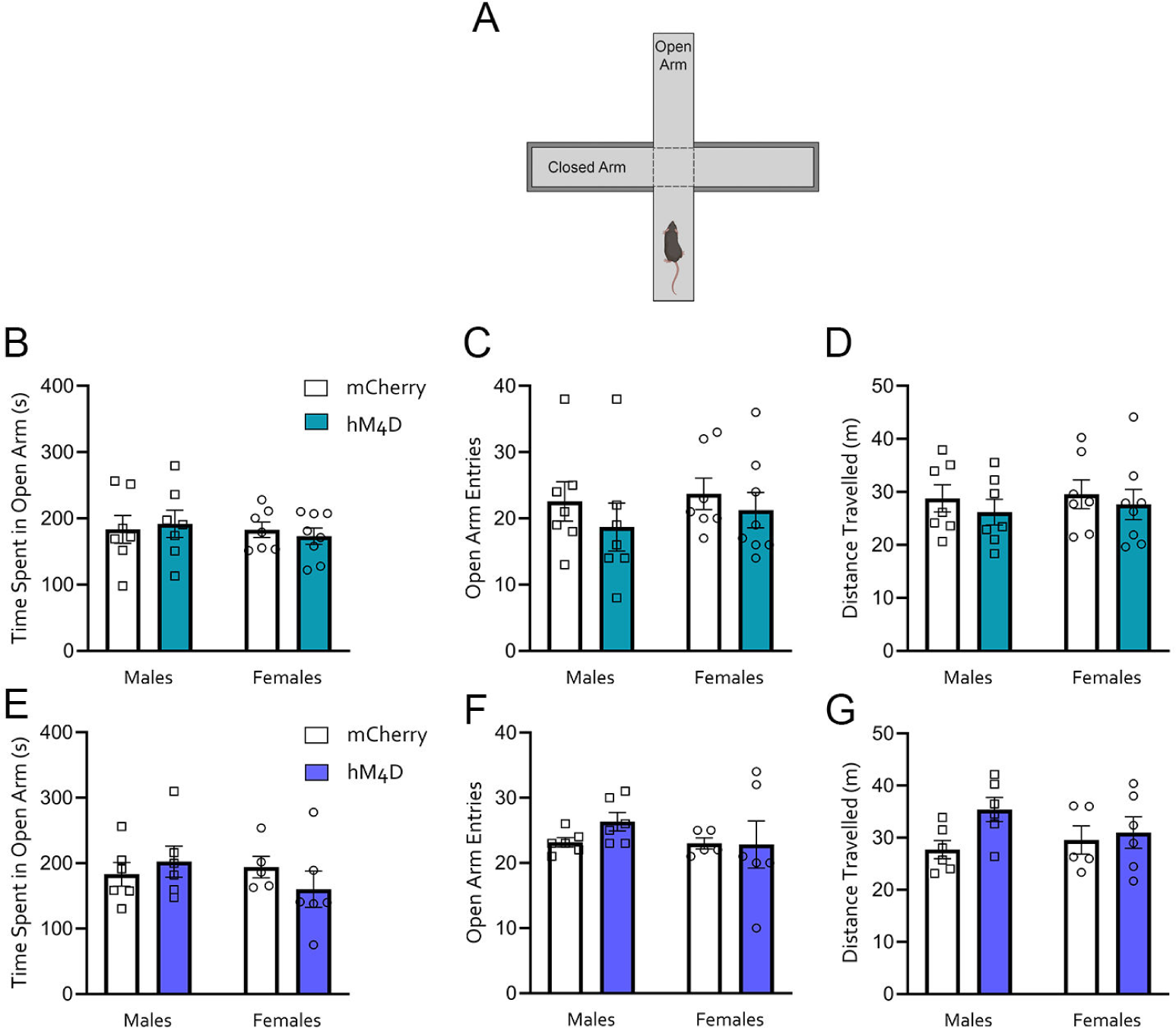
Chronic inhibition of the vCA1-mPFC pathway does not affect long-term performance in the elevated plus maze. **A.** Following the experiments from Figure 6, animals in both the juvenile and adult cohorts were tested in the elevated plus maze (EPM) in the absence of C21. **B-D**. Juvenile cohort. mCherry and hM4D animals that underwent C21 treatment from P15-P30 did not differ in time spent in the open arms of the EPM (**B**), number of open arm entries (**C**) or distance travelled (**D**). mCherry n= 14 (7 females, 7 males); hM4D n= 15 (8 females, 7 males). **E-G**. Adult cohort. mCherry and hM4D animals that underwent C21 treatment from P60-P75 showed equivalent time spent in the open arms of the EPM (**E**), number of open arm entries (**F**) or distance travelled (**G**). mCherry n= 11 (5 females, 6 males); hM4D n= 12 (6 females, 6 males). Individual female datapoints are represented as circles, and males as squares.

## Notes

**Declaration of interests:** The authors declare no competing interests.

### Competing Interest Statement

The authors have declared no competing interest.

### Summary of Updates

All figures were revised to incorporate larger n, a new chronic manipulation of iCA1-mPFC pathway in juveniles and adults (Fig 6J-Q), as well as characterization of the effects of vCA1- and iCA1-mPFC chronic manipulations on mPFC spontaneous transmission (Fig 7).

